# Seeing the future: predictive control in neural models of ocular accommodation

**DOI:** 10.1101/2021.12.09.471909

**Authors:** Jenny C. A. Read, Christos Kaspiris-Rousellis, Toby S. Wood, Bing Wu, Björn N.S. Vlaskamp, Clifton M. Schor

## Abstract

Ocular accommodation is the process of adjusting the eye’s crystalline lens so as to bring the retinal image into sharp focus. The major stimulus to accommodation is therefore retinal defocus, and in essence, the job of accommodative control is to send a signal to the ciliary muscle which will minimize the magnitude of defocus. In this paper, we first provide a tutorial introduction to control theory to aid vision scientists without this background. We then present a unified model of accommodative control that explains properties of the accommodative response for a wide range of accommodative stimuli. Following previous work, we conclude that most aspects of accommodation are well explained by dual integral control, with a “fast” or “phasic” integrator enabling response to rapid changes in demand, which hands over control to a “slow” or “tonic” integrator which maintains the response to steady demand. Control is complicated by the sensorimotor latencies within the system, which delay both information about defocus and the accommodation changes made in response, and by the sluggish response of the motor plant. These can be overcome by incorporating a Smith predictor, whereby the system predicts the delayed sensory consequences of its own motor actions. For the first time, we show that critically-damped dual integral control with a Smith predictor accounts for adaptation effects as well as for the gain and phase for sinusoidal oscillations in demand. In addition, we propose a novel proportional-control signal to account for the power spectrum of accommodative microfluctuations during steady fixation, which may be important in hunting for optimal focus, and for the nonlinear resonance observed for low-amplitude, high-frequency input. Complete Matlab/Simulink code implementing the model is provided at https://doi.org/10.25405/data.ncl.14945550.

## Introduction

Accommodation refers to the ability of the eye to change its focus between near and far distances, so as to ensure that images remain in sharp focus at the fovea across a wide range of object distances. This is achieved by changes in the convexity of the intra-ocular lens, brought about by contraction of the ciliary muscle (Figure 1). To focus on distant objects, the ciliary muscle is relaxed, the lens curvature and thus its optical power is minimal; to focus on near objects, the ciliary muscle contracts, the lens curvature increases and so does its optical power. Accommodation is usually controlled by the brain as an unconscious reflexive process.

**Figure 1.**
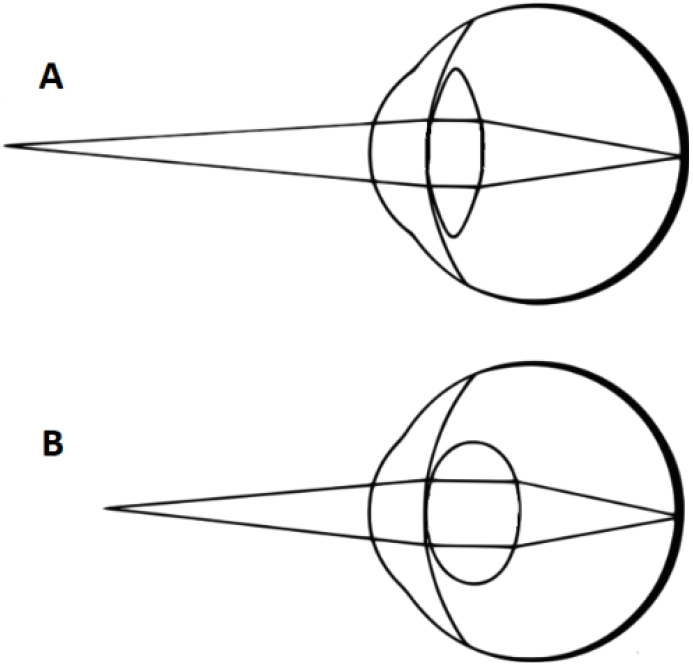
(A) Accommodating on a distant object. When the ciliary muscle is slack, tension in the suspensory zonules is released and the intra-ocular crystalline lens flattens, enabling distant objects to appear in focus on the retina (for an emmetropic eye). Light from a nearby object, such as shown, is therefore out of focus. (B) Accommodating on a nearby object. The ciliary muscle has contracted, increasing the curvature of the lens (blue arrows) in order to bring nearby objects into focus. Not to scale. Image: Pearson Scott Foresman, public domain.

A full understanding of this process requires a knowledge of (i) the optical and biomechanical properties of the eye; (ii) how the required accommodative response is derived from retinal and extra-retinal cues; and (iii) the neural signals controlling the ciliary muscle to bring about this response. In this paper, we concentrate on the third of these.

We begin by discussing the basic structure of models of neural control of accommodation. A key goal of this section is to provide a clear review of the subject, introducing concepts and summarizing previous work in a way which is accessible to vision scientists without a background in classical control theory. Accordingly, this section incorporates a tutorial to bring such readers up to speed.

The core of accommodative control is a negative feedback loop attempting to null the error between accommodative demand, i.e. the accommodation at which the fixated object will be in sharp focus, and response, i.e. the accommodation actually adopted. Such feedback loops are vulnerable to instabilities caused by the finite latencies within the control system. A well-established strategy for avoiding such instabilities is to predict the eye’s response to a motor command. This requires an “efference copy” of the signal sent to the ocular plant, along with an internal or “forward” model of the plant, enabling the system to predict the response to the motor signal. Control can then be based on the predicted future input, rather than the currently sensed input, effectively removing the effect of the latencies. We consider the particular form known as a Smith predictor (Abe & Yamanaka, 2003; Miall et al., 1993; Smith, 1957), designed for closed-loop control of systems with long delays in the feedback. Predictive models stand in contrast to classical models which do not take account of the sensory consequences of the body’s own motor actions.

Armed with this background, we next discuss the evidence that accommodation uses a Smith predictor, and examine empirical constraints on the model parameters. We aim to produce a model which can account for behavior in both steady-state and smooth tracking, including accommodative lag/lead, adaptation, critical damping, and Bode plots of gain and phase. (Extending the model to reproduce dynamics of the step response (Bharadwaj & Schor, 2005, 2006; Schor & Bharadwaj, 2004, 2006) will be covered in a subsequent paper.)

This analysis leads us to conclude that accommodative control most likely incorporates a predictor, in order to avoid instabilities due to the sensorimotor latency. By “predictor”, we mean a forward model to predict the effect of commanded accommodation changes on the visual input. The evidence that the system predicts changes in stimulus demand is equivocal, and our model simply assumes that demand does not change over the timescale of the latency.

We conclude that accommodation can be modelled successfully as a predictive system with integral control, but that there are fairly tight constraints on the gain and time-constant of the integral controller in order for the system to be consistent with empirical data for step and smooth tracking. Following previous work, we add a slow, second-order integral controller to account for adaptation effects, and show that care is required when using this dual-control with predictive models.

Most accommodation models omit noise, but noise provides important constraints on model structure and parameters. Predictive models can end up amplifying internal noise when the defocus signal is removed e.g. by viewing through pinholes, which is not observed empirically. Avoiding these resonances places further constraints on model parameters. An important contribution of this paper is that our model explicitly includes noise.

Noise also accounts naturally for the fluctuations seen in steady-state accommodation. These are often called microfluctuations although they are actually quite substantial at around ±0.5D, exceeding the depth of field (Campbell et al., 1959a; Charman & Heron, 1988, 2015; Kotulak & Schor, 1986b). The source and purpose of these is unclear: as well as noise, they may reflect disturbances from the intraocular pulse, mechanical resonances within the ocular plant, deliberate attempts at “hunting” in order to find the best point of focus, and/or fluctuating input from other influences on accommodation (Charman & Heron, 1988, 2015; Collins et al., 1995; Denieul, 1982; Gray et al., 1993b).

In normal viewing, the power spectrum of microfluctuations typically shows a pronounced peak at around 2Hz. This peak is much weaker when viewing through pinhole pupils, where link between accommodation and image quality is cut (“open-loop”). This may be because in bright viewing conditions, where the pupil stops down and depth of focus is large, microfluctuations are of no assistance in improving vision, and might even cause ocular fatigue. In our model, we are able to reproduce this behavior by including an additional control signal which is driven directly by sensed defocus and *not* by the output of the Smith predictor.

Putting these different components together results in a model where accommodation is controlled by the sum of four separate neural signals. The model has a total of ten parameters (Table 2), most of which are quite tightly constrained by the data. In the Results section, we present simulations demonstrating that this model can account simultaneously for a wide range of observations.

## Methods

### Accommodation as a linear, time-invariant negative feedback control system

> *“A complex system that works is invariably found to have evolved from a simple system that worked. A complex system designed from scratch never works and cannot be patched up to make it work. You have to start over with a working simple system.” – Gall’s Law* (Gall, 1977).

In the spirit of Gall’s Law, we begin with the simplest possible conceptual model of accommodation (Figure 2). Viewed as a whole, the model has one input, *accommodative demand*, corresponding to the vergence of light rays from the object we wish to look at. This is measured in diopters; the demand in diopters corresponds to the reciprocal of the distance in metres from the eye. For an infinitely far object, the demand is 0D; for an object at 50cm, the demand is 2D.

**Figure 2.**
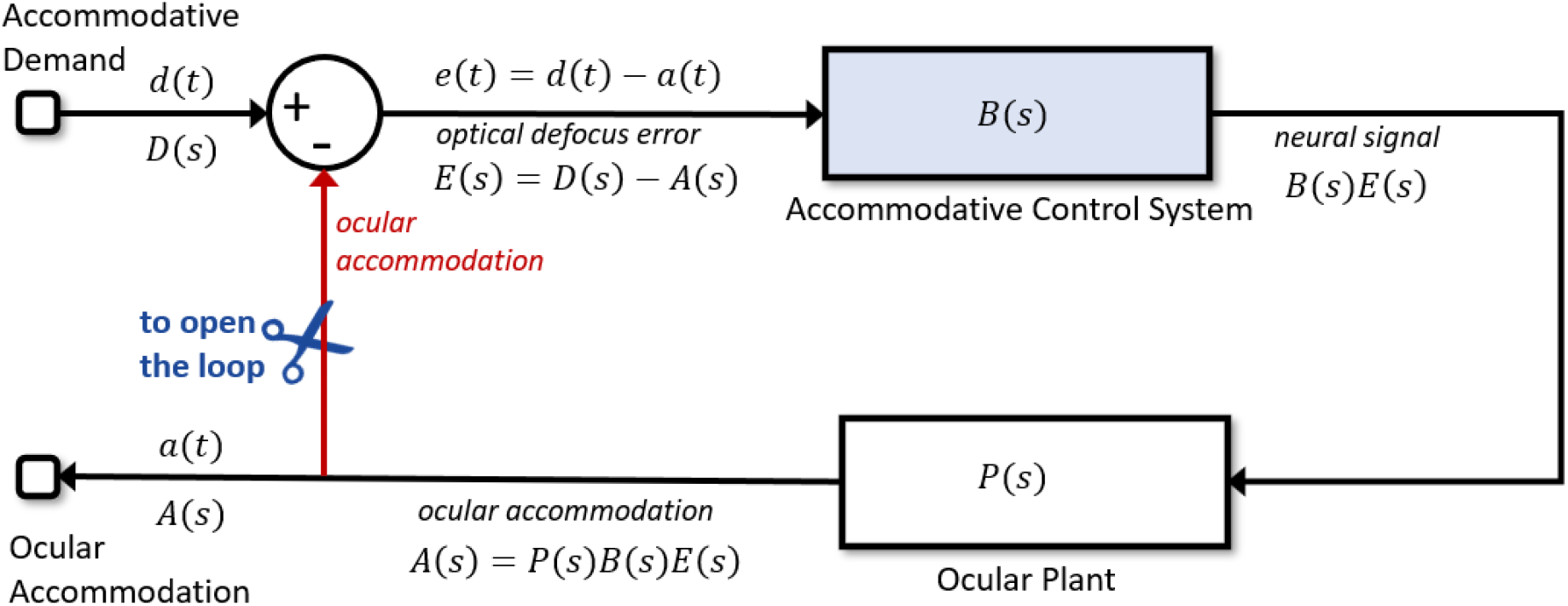
Conceptual model of accommodation. There is a feedback loop, whereby the output (accommodation) affects the input to the control system. The blocks labelled Accommodative Control System and Ocular Plant are shown here as “black boxes” which take inputs and yield outputs, without showing how the output is computed. Their transfer functions are B(s) and P(s) respectively. The input to the overall system is the accommodative demand, reflecting the distance of the fixated object, and the output is the ocular accommodation, i.e. where the eye is focused. Defocus error is the difference between these, demand minus accommodation. Signals are shown in the time-domain, e.g. d(t), and as Laplace transforms, e.g. D(s). When the system is driven in “open-loop” mode, the connection shown in red is effectively severed at the scissors icon, so that the input to the Accommodative Control system becomes independent of ocular accommodation.

The model also has one output, *ocular accommodation*. When the eye is correctly accommodated, the accommodation will be equal to the demand so that the image is in focus on the posterior receptor layer of the retina. *Defocus* is the difference between the accommodative demand and the ocular accommodation, all measured in diopters. It acts as an error signal to the model. As discussed in the Introduction, we assume that defocus is a single, signed value which is somehow computed by the visual system from the retinal image (e.g. using blur, higher-order aberrations, longitudinal chromatic aberration (Burge & Geisler, 2011; Cholewiak et al., 2018; Fincham, 1951; Kruger et al., 1993; Seidemann & Schaeffel, 2002; Wilson et al., 2002)) and represented as a neural error signal; how this is achieved is beyond the scope of this paper. In our sign convention, positive defocus error means that the eye is not accommodating enough, i.e. the eye is focusing on a point more distant than the object of interest, so the ocular image is focused behind the retina. Positive defocus error should therefore stimulate an increase of accommodation. The accommodative control system takes the defocus error as input and uses it to compute a neural control signal (blue block in Figure 2). This neural signal is then fed into the ocular plant block in Figure 2. This block, corresponding physiologically to the ciliary muscle, lens and other components, converts the neural signal into the optical power of the lens, i.e. the ocular accommodation. This in turn affects the defocus error, since defocus is demand minus accommodation. The accommodative control system is designed to adjust accommodation so as to minimise the defocus error signal (Toates, 1972). Thus, this is a negative feedback system.

In any negative feedback system, one faces the question of how to choose the control signal so as to minimize the error. One obvious form of error correction is to make the corrective signal proportional to the error. For example, a very simple form of automotive cruise control might make acceleration proportional to the difference between the current and the desired speed. Other widely-used possibilities are to integrate the error over time, or to anticipate changes by including a term scaling with the derivative. Together, control systems of this type are called PID (proportional-integral-derivative) controllers.

In reality, of course, defocus is not the only visual cue to accommodation (Heath, 1956b; Maddox, 1893). One additional component that we discuss below and include in our models is the system’s bias towards a particular baseline or resting accommodation (see (Rosenfield et al., 1993a) for a review). Factors which for simplicity we neglect in this paper include pictorial cues to distance, sensed proximity and crosslinks from the vergence system. However, defocus is the only visual cue which is itself altered by accommodation, and thus the cue intrinsic to the negative feedback loop.

#### Modelling neural signals as if they were in diopters

In this initial part of the paper we will keep the discussion as general as possible, without committing to a particular model of the Ocular Plant or Accommodative Control System blocks shown in Figure 2. However, one detail is worth noting. Without loss of generality we will set the overall gain of the plant to 1, meaning that it passes a constant signal unchanged. In reality, the neural signal is encoded in spikes per second, and the output of the ocular plant is accommodation in diopters. There must therefore be a gain or conversion factor within the neural signal which converts spikes per second into diopters, taking into account the biomechanical gain of the plant (Gamlin et al., 1994). Without loss of generality, we can fold this conversion factor into our neural signals. Thus by setting the plant gain to 1, we represent all the neural signals in the model as if they were diopters. This makes them particularly simple to interpret.

#### Closed-loop versus open-loop

The model shown in Figure 2 is “closed-loop”: that is, the input to the accommodative control system (defocus error) is affected by its output (ocular accommodation). As discussed, this forms a negative feedback loop, in which increases in defocus error stimulate changes in accommodation that in turn reduce defocus error.

If we use the scissors shown in Figure 2 to cut the connection shown in red, we obtain the equivalent open-loop system, in which the output of the system has no effect on its defocus error. It might seem impossible to cut the connection in this way in the living eye, but in fact all that is required to examine the open-loop mode is to make the optical error signal independent of the accommodative response. There are two main ways in which this can be done. First, by measuring accommodation and optically adding the current accommodation state onto the current input demand. The eye’s own optics then effectively remove accommodation, so that the error signal forming the input to the visual system is simply the demand applied by the experimenter, independent of the accommodative response. A positive non-zero open-loop error signal continues to stimulate increases of the accommodation response until it reaches saturation, reminiscent of a dog chasing its tail.

Alternatively, the optical error signal can be set to zero by using a pinhole pupil. Through small pinholes, objects appear slightly blurred due to diffraction, but critically, this blur is virtually independent of the stimulus accommodative demand or the ocular accommodation. Pinholes do not cause a “dog chasing tail” accommodative response; rather accommodation tends to assume its resting state. This suggests that the visual system experiences images seen through pinholes as having zero defocus. Thus, viewing through pinholes is a special case of open-loop in which the input is effectively clamped to zero regardless of output. As we shall see, examining a system in open-loop mode can produce valuable information about its function.

#### Primer on control system theory

At this point, we note that vision scientists may not be familiar with the classical control systems approach taken in this paper. This section aims to provide a bare-bones introduction to enable such readers to follow subsequent sections. Table 1 provides a reference for all the symbols used throughout the paper.

**Table 1.** Symbols used in this paper.

| Symbol | Meaning |
| --- | --- |
| $\omega$ | Angular temporal frequency, $\omega = 2\pi f$ |
| $\phi(f)$ | phase-delay of accommodation at frequency $f$ |
| $\tau_{\text{fast}}$ | Time-constant of the fast leaky-integrator controller, see Equation 10 |
| $\tau_{\text{plant}}$ | Time-constant of the ocular plant, when this is modelled as a leaky integrator, see Equation 9 |
| $\tau_{\text{slow}}$ | Time-constant of the slow leaky-integrator controller, see Equation 12 |
| $\zeta$ | Damping coefficient, see Equation 20 |
| $\hat{a}(t + T_{\text{mot}})$ | Predicted accommodation at a time $T_{\text{mot}}$ after the current time $t$. In this paper, generally assumed equal to $a(t+T_{\text{mot}})$, i.e. prediction is perfect. |
| $\hat{d}(t + T_{\text{mot}})$ | Predicted demand at a time $T_{\text{mot}}$ after the current time $t$. In the no-change prediction model, this is assumed to be the same as the last available demand, from time $T_{\text{sens}}$ before the current time, i.e. $d(t-T_{\text{sens}})$ |
| $A(s)$ | Laplace transform of accommodation relative to rest focus |
| $a_{\text{RF}}$ | Rest focus, i.e. accommodation adopted in the absence of any visual stimulus |
| $a_{\text{ss}}$ | Steady-state accommodation in response to $d_{\text{ss}}$, see Equation 7 |
| $C(s)$ | Transfer function of controller |
| $D(s)$ | Laplace transform of accommodative demand relative to rest focus |
| $d_{\text{ss}}$ | Steady-state demand, see Equation 7 |
| $E(s)$ | Laplace transform of defocus error, $E(s)=D(s)-A(s)$ |
| $f$ | Temporal frequency |
| $g(f)$ | gain of accommodation at frequency $f$ |
| $G_{\text{fast}}$ | Steady-state gain of the fast leaky-integrator controller, see Equation 10 |
| $G_{\text{open}}$ | steady-state open-loop gain of accommodation |
| $G_{\text{slow}}$ | Steady-state gain of the slow leaky-integrator controller, see Equation 12 |
| $H_{\text{closed}}(s)$ | Closed-loop transfer function relating demand to accommodation, see Equation 4 |
| $H_{\text{open}}(s)$ | Open-loop transfer function relating demand to accommodation, see Equation 3 |
| $j$ | Square root of -1. |
| $P(s)$ | Transfer function of ocular plant |
| $s$ | Complex temporal frequency in Laplace domain, $s = j\omega$, see Equation 1 |
| $t$ | Time |
| $T_{\text{lat}}$ | Total sensorimotor latency, $T_{\text{lat}} = T_{\text{sens}} + T_{\text{mot}}$ |
| $T_{\text{mot}}$ | Motor latency, i.e. time taken for the neural signal controlling accommodation to travel from the brain to the ocular plant |
| $T_{\text{sens}}$ | Sensory latency, i.e. time taken for defocus at the retina to reach the accommodative control system in the brain |

**Table 2.** Parameter values for the Simulink model supplied with the paper and used to obtain the results (except where noted otherwise in figure legends). These values are visible in the Simulink Model Workspace, and can be altered there if desired.

| Parameter | Symbol used in the paper | Name in Simulink workspace | Value |
| --- | --- | --- | --- |
| Rest focus | $a_{RF}$ | RestFocus | 1.4D |
| Sensory latency | $T_{sens}$ | SensoryLatency | 0.20s |
| Motor latency | $T_{mot}$ | MotorLatency | 0.10s |
| Time constant of plant | $\tau_{plant}$ | PlantTimeConstant | 0.156s |
| Gain of fast integrator | $G_{fast}$ | FastGain | 8.0 |
| Time constant of fast integrator | $\tau_{fast}$ | FastTimeConstant | 2.5s |
| Gain of slow integrator | $G_{slow}$ | SlowGain | 5.0 |
| Time constant of slow integrator | $\tau_{slow}$ | SlowTimeConstant | 100s |
| Noise power |  | NoisePower | 0.001 with sample time 0.01s |
| Where to clip the proportional signal |  | ProportionalClipping | 0.15D |

##### Linear time-invariant (LTI) systems and the Laplace domain

Linear systems are those whose outputs for a linear combination of inputs are the same as a linear combination of individual responses to those inputs. For example, in Figure 2, if the system were linear, then if demand timecourse *d*_1_(t) elicited accommodation response *a*_1_(t), and demand *d*_2_(t) elicited *a*_2_(t), the response to a new demand made up of a weighted sum of these two timecourses, *w*_1_*d*_1_(t)+*w*_2_*d*_2_(t) would be *w*_1_*a*_1_(t)+*w*_2_*a*_2_(t). A time-invariant system is one where the same input, delayed by a time *T*, will always elicit the same response, also delayed by a time *T*. Thus if demand *d_1_*(*t*) elicited accommodation response *a*_1_(*t*), demand *d*_1_(*t*-*T*) would elicit accommodation response *a*_1_(*t-T*).

Where a system is both linear and time-invariant (LTI), its response can be analysed using Laplace transforms of the variables. The Laplace transform turns integral and differential equations into polynomial equations which are much easier to solve. Time-domain functions are converted into Laplace-domain functions of a complex frequency variable *s*. We assume that all signals are zero for times before t=0, and write the Laplace transform of a signal *f(t*) as *F*(*s*), where

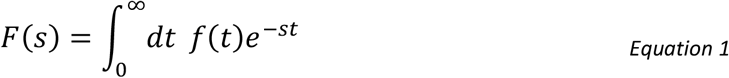

We will adopt the convention where, when a lower-case variable represents a function of time *t*, the corresponding upper-case denotes its Laplace transform as a function of *s*. The Laplace transform is closely related to the Fourier transform with which vision scientists are typically more familiar, with *s* representing a complex version of angular temporal frequency: *s=jω* (where we use *j* for the square root of −1 throughout).

In a circuit diagram like Figure 2, the effect of an LTI block is simply to reweight the amplitude, and/or shift the phase, of each frequency in the input. This means that each LTI block can be written simply in terms of its complex *transfer function H*(*s*). As discussed in more detail below, a transfer function *H*(*s*) is a kind of gain, since it is the ratio of the output to the input, for each frequency *s*. For example, consider a transport delay block, whose effect is to delay the input signal by a latency *T*, and which thus shifts the phase of each frequency. If the input signal is *i(t)*, the output after delay is *o(t)* = *i(t-T)*. Substituting this into Equation 1, we find that

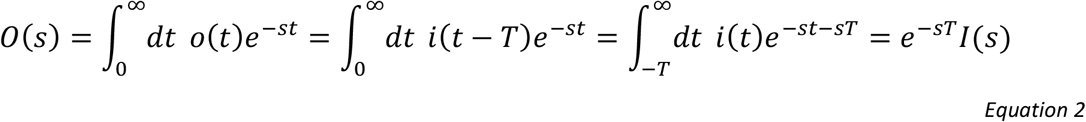

where we used the fact that *i*(*t*)=0 for *t*<0. Thus, the transfer function of a transport delay block is *H*(*s*)=exp(−*sT*). Constant signals are unaffected (H(0)=1); time-varying signals undergo a shift in phase proportional to their temporal frequency.

Integrating by parts, and using the assumption that *f*(0)=0, we see that

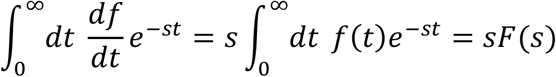

and so the Laplace transform of a derivative is just *s* times the Laplace transform of the original function. This means that differentiation can be represented very simply in Laplace space by multiplication by *s*, and integration by 1/*s*.

In LTI systems, one can do algebra on the Laplace transforms in the usual way. The transfer function for several LTI systems in parallel is the sum of the individual transfer functions, while the transfer function for several LTI systems in series is the product of the transfer functions for the individual systems.

##### A mathematical trick to handle rest focus

When viewing through pinholes, although the demand is zero, accommodation tends not to be zero but to converge on a “rest focus”, *a*_RF_, generally of around 1.4D (Leibowitz & Owens, 1978; Rosenfield et al., 1993b), which is the value we shall assume for our model. A similar default focus is also observed in darkness. To account for this, we assume that the accommodative control system adds onto the signal computed from defocus a constant “bias” signal. Because we have normalized neural signals to be expressed in diopters, setting this bias signal equal to the rest focus ensures that accommodation returns to the rest focus if the defocus error is clamped at zero.

This bias signal leads to a small complication, because it technically violates the assumption that all signals are zero for *t*≤0. To handle this, we express both accommodation and demand relative to the rest focus. We define *A*(*s*) to be the Laplace transform, not of accommodation itself, but of accommodation relative to rest focus, *a(t)-a_RF_*. Similarly *D*(*s*) is the Laplace transform of demand relative to rest focus, *d(t)-a_RF_*. With this trick, we can then analyse the system in the Laplace domain as if there were no bias signal (*a*_RF_=0), and at the end simply add *a*_RF_ back on to demand and accommodation when we move back to the time domain. All the analyses in this paper use this approach.

##### Open- and closed-loop transfer functions

Where accommodation is driven in open-loop mode (imagine Figure 2 after the scissors have cut), we have

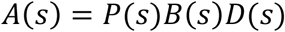

where *B(s)* is the transfer function representing the brain’s accommodative control system and *P(s)* that representing the ocular plant. As described in the previous section, *A*(s) and *D*(s) are the Laplace transforms of accommodation and demand relative to rest focus. The open-loop transfer function relating output *A*(s) (accommodation) to input *D*(s) (demand) is thus

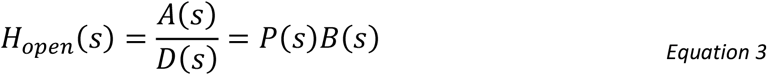

In closed-loop mode (Figure 2 with no scissors), the input to the accommodative control system is defocus error, *E*(s) = *D*(s)-*A*(s). We therefore now have

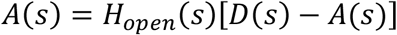

and thus derive the closed-loop transfer function:

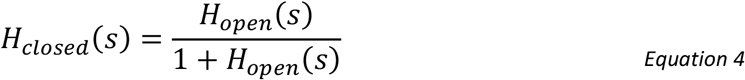

where

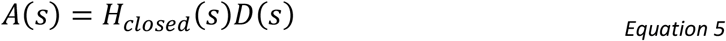

This relationship between the open- and closed-loop transfer functions is a standard result for a feedback loop like the one in Figure 2.

##### Steady-state response, accommodative lag and lead

LTI theory shows that the steady-state response is obtained by evaluating the system at *s*=0 (zero frequency). So, if we apply a constant demand *d*_ss_ in closed-loop mode, Equation 5 becomes

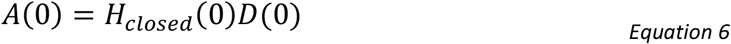

where *D*(0)=*d*_ss_-*a*_RF_ and A(0)=*a*_ss_-*a*_RF_ (recalling that accommodation and demand are defined relative to rest focus *a*_RF_). From Equation 4, we can write *H_closed_*(0) in terms of *H_open_*(0). It will be convenient to introduce the notation *G*_open_ for *H*_open_(0), i.e. the open-loop steady-state gain of the system. Putting this together with Equation 4 and Equation 6, we find that accommodation will eventually be

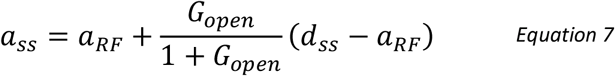

The steady-state defocus error is

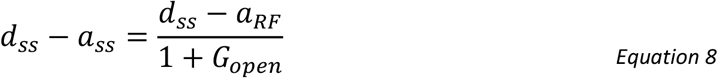

Equation 8 shows that – regardless of the control system or plant – the defocus error will be zero when the demand is equal to the rest focus. This is natural enough, since the rest focus is the value to which the system is biased.

However, for other demands, the steady-state error is not zero. When the demand is nearer than the rest focus, the accommodative response remains further than the demand, a situation referred to as accommodative lag. Conversely when demand is further than rest, accommodation is nearer than demand; this is accommodative lead.

Importantly, the amount of the error depends on the steady-state open-loop gain *G*_open_. This demonstrates an important property of negative-feedback systems which attempt to minimise error: small error requires high open-loop gain. Since we have set the gain of the plant to 1 (without loss of generality, as noted above), the gain *G*_open_ is set entirely by the brain’s accommodative control system. Empirically, accommodation reaches around 80%-90% of the demand when the demand is far from the rest focus. From Equation 4, we have

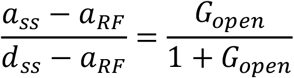

so the observation that accommodation is around 80-90% of demand implies that *G*_open_/(1+*G*_open_) is around 0.8-0.9, and in turn that Gopen must be in the range 4-9.

##### Gain and phase of response to sinusoidal inputs

A property of any LTI system is that (after initial onset transients have died away) its response to a sinusoidal input is a sinusoidal output, with a gain and phase reflecting the transfer function of the system. Specifically, if the closed-loop transfer function is *H_closed_*(*s*), then if accommodative demand is a sinusoidal function of time, the accommodative response will also be a sinusoid with the same temporal frequency *f*. The amplitude of the response will be the amplitude of the demand multiplied by the gain at that frequency, *g*(*f*), and the phase will be delayed by *ϕ*(*f*). We will use lower-case *g*(*f*) to denote the gain of a system at a particular temporal frequency *f*, and upper-case *G*=*g*(0) to denote the steady-state gain, as we did above for *G*_open_. According to a standard result of LTI theory, the gain and phase-delay of an LTI system at frequency *f* can be obtained from the complex number represented by its transfer function *H*(*s*) evaluated at *s*=2π*jf*. The gain *g(f)* is the magnitude of the complex number *H*(2π*jf*) and the phase-delay *ϕ*(*f*) is its phase.

Sometimes below for brevity we will refer to “the gain” of an LTI operator, without specifying a frequency. In this case, we mean its steady-state gain. For example, when we refer to “the gain” of a low-pass filter, we mean the ratio of its steady-state output to a constant input.

### Sensorimotor latencies: a problem for control

These preliminaries out of the way, we now consider different possibilities for the contents of the blue block labelled Accommodative Control System in Figure 2. We begin by expanding this block as shown in Figure 3. We now explicitly include the rest focus signal discussed above. But critically, Figure 3 now also shows the system’s latency, which we have divided into two parts. The first is an afferent-sensory latency, representing the time taken for information about the retinal image to travel up the optic nerve and for the brain to compute a signed estimate of defocus, for example using longitudinal chromatic aberration or higher-order aberrations. The second is an efferent-motor latency, representing the time taken for the resultant neural signal to travel from the Edinger-Westphal nucleus down the IIIrd cranial nerve, relay in the ciliary ganglion and reach the ciliary muscle. The motor latency is reduced by the fact that the axons from the ciliary ganglion to the ciliary muscle are myelinated, unusually for postganglionic axons of the autonomous nervous system (Tamm & Lütjen-Drecoll, 1996; Warwick, 1954). The sensory and motor latencies have been estimated as *T*_sens_~200ms and *T*_mot_~100ms respectively (Gamlin et al., 1994; Schor et al., 1999; D. Wilson, 1973), and we will fix the values in our model at these values. In Figure 3, these latencies are shown within the Accommodative Control System, i.e. the brain, but the model functioning is unchanged if, for example, part of the motor latency occurs at a neuromuscular junction in the eye, or indeed if both latencies are merged into a single block.

**Figure 3.**
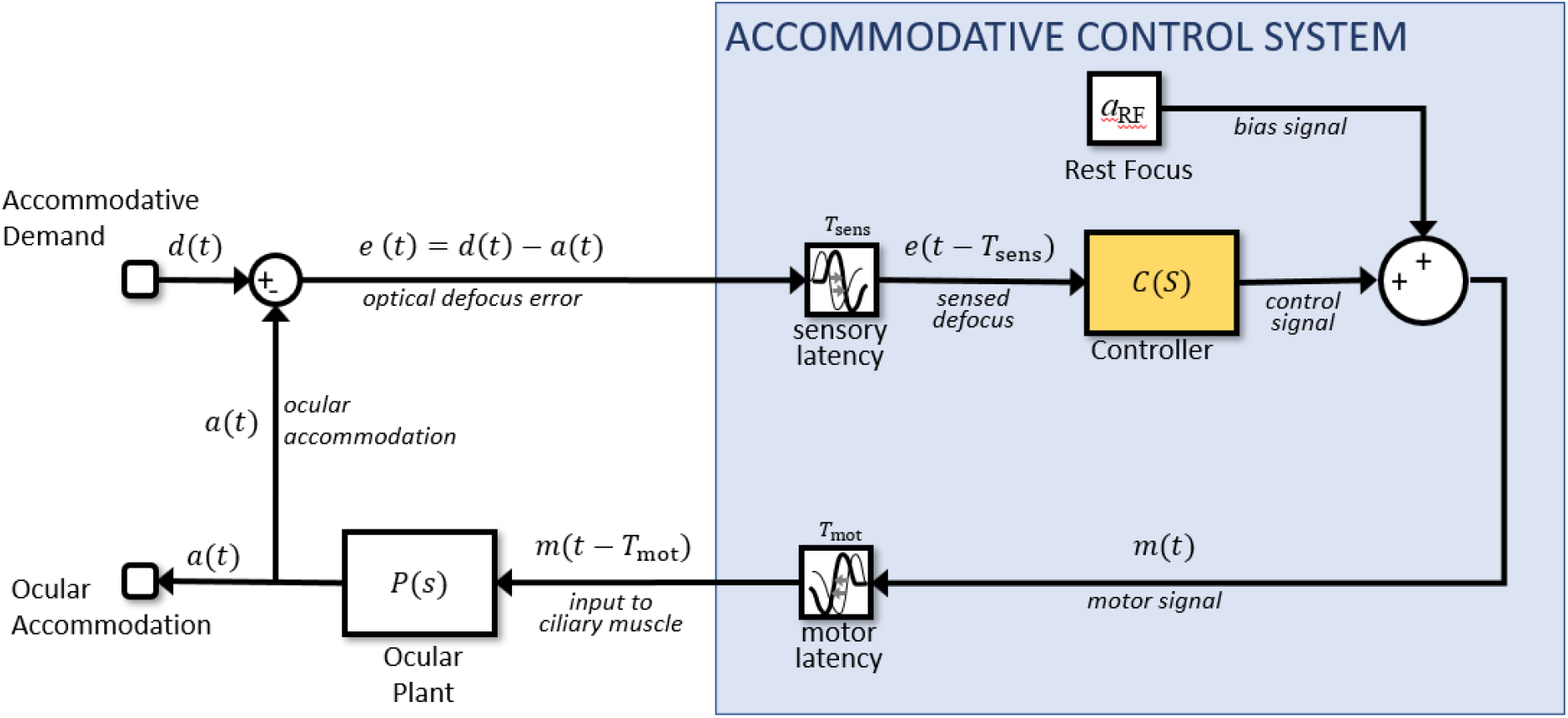
Expanding the conceptual model shown in Figure 2 so as to show the rest focus and sensorimotor latencies. This is the same circuit diagram, but the block labelled Accommodative Control System has here been expanded to explicitly show the constant bias signal accounting for the rest focus, and the latencies. There is a sensory latency T_sens_ before the retinal defocus signal reaches the controller, and a further motor latency T_mot_ before the neural signal reaches the plant.

Latencies are a potentially serious problem for any control system. In the block diagram shown in Figure 3, we can see that the defocus error only becomes available to the block marked Controller after the sensory latency. The controller therefore operates not on *e*(*t*), but *e*(*t-T*_sens_): the retinal defocus as it was a time *T*_sens_ ago. This in turn reflects the accommodation due to the neural signal sent up to a time *T*_sens_+*T*_mot_ ago. Thus, the system suffers an overall latency of *T*_lat_ = *T*_sens_ +*T*_mot_. This can easily lead to overshoots and “ringing”: oscillations in accommodation as the system is driven beyond the correct value by the out-of-date error signal.

Overshoots and ringing due to an out-of-date error signal would be seen with the response to step changes in demand, but in fact the second-order dynamics already indicate that LTI models do not suffice to account for the response to large step changes; accommodative control seems to have special mechanisms for these which are beyond the scope of this paper (Bharadwaj & Schor, 2005, 2006; Schor & Bharadwaj, 2004, 2005). However, an out-of-date error signal would also affect the response to sinusoidal oscillations in demand which we will concentrate on in this paper.

Empirically, accommodation shows a low-pass response: gain is greatest in the steady-state, and decreases monotonically with temporal frequency (Charman & Heron, 2000; Krishnan et al., 1973; Kruger & Pola, 1986; Ohtsuka & Sawa, 1997; Stark et al., 1965). However, it is challenging to achieve this with the circuit diagram shown in Figure 3 and a Controller block which is simply a PID controller. Because of the latency, the system can easily end up out of phase, so that the changes in accommodation actually enhance the defocus rather than reducing it, as intended. This shows up as resonances or local peaks in the gain function, making it non-monotonic. This is not observed empirically.

#### Overcoming latencies with a predictive control system: the Smith Predictor

The solution seems to be that the visual system actually bases its neural control not on the currently available sensed value of retinal defocus, but on its internal prediction of the *future* retinal defocus. That is, whereas in Figure 3 the controller operates on the sensed defocus, which due to the sensory latency actually represents defocus as it was some time in the past, in a predictive model the controller operates on the predicted *future* defocus (Smith, 1957). Figure 4 shows how Figure 3 can be modified so that the input to the controller is predicted future defocus. Defocus is the difference between the stimulus accommodative demand and the ocular accommodation, so predicting future defocus requires a prediction both of demand and accommodation.

**Figure 4.**
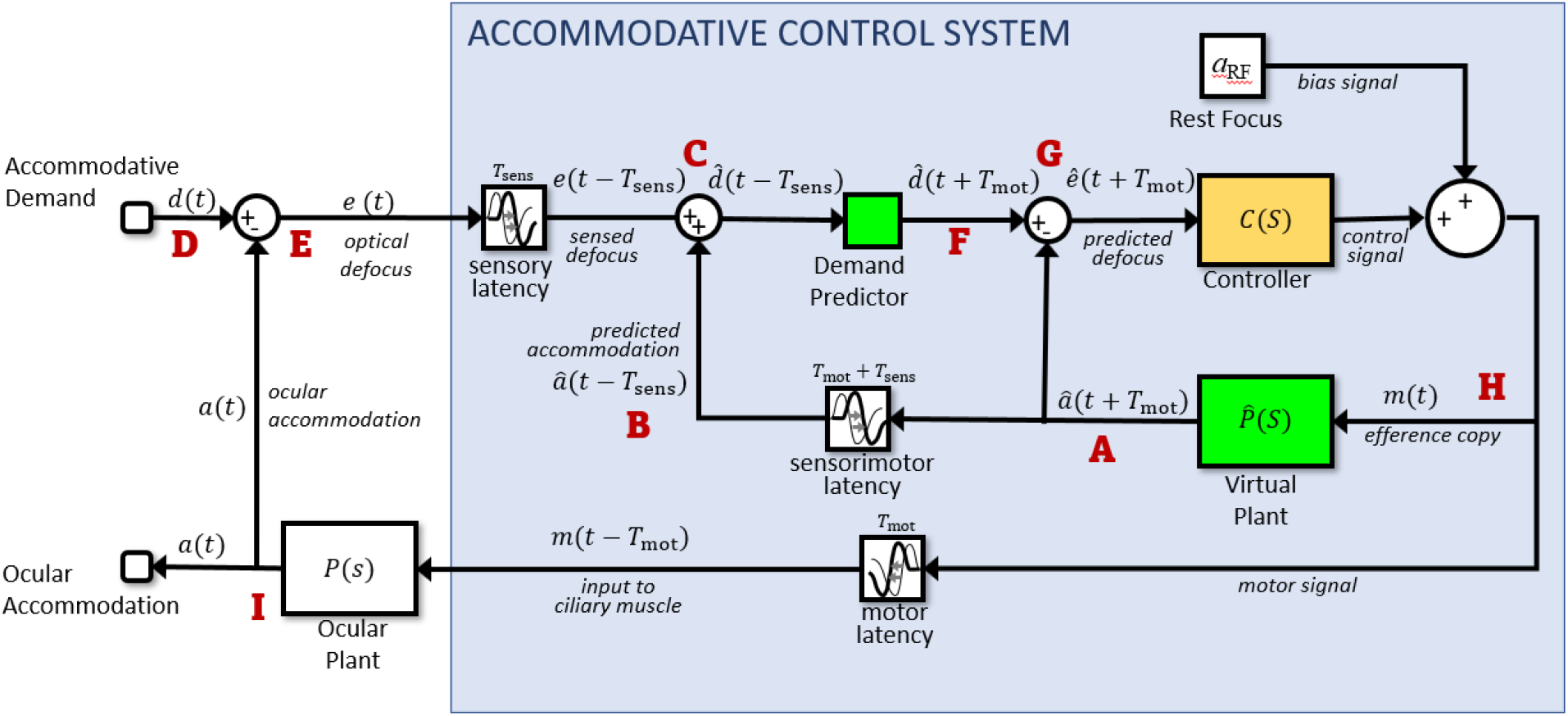
Predictive control. Compare with Figure 3: the Controller block has been replaced with a more complex system including two predictive blocks (green) as well as the original Controller block (yellow). The prediction helps avoid instability due to the sensorimotor latencies. To predict accommodation, the model includes a Virtual Plant block (forward model) to compute what accommodation will be a time T_mot_ in the future, i.e. after the motor latency. If the forward model is accurate, this can in principle predict accommodation perfectly up to t+T_mot_, since accommodation is under the system’s own control. To predict demand at time T_mot_ into the future, the model uses a Demand Predictor block. This requires extrapolating demand at time T_lat_=T_sens_+T_mot_ beyond the last available estimate. This is unlikely to be entirely accurate, since demand can reflect changes in the outside world, beyond the system’s control. Red labels indicate locations referred to in the text.

The brain is in principle able to predict accommodation perfectly up to future times less than the motor latency, simply based on the signals it has already sent to the accommodative plant. (Campbell & Westheimer, 1960; Hung et al., 2002; Krishnan et al., 1973; Schor & Bharadwaj, 2004; Stark et al., 1965; Sun et al., 1989) To do this, the visual system must effectively have its own internal model of the ocular plant, represented by the *Virtual Plant* block in Figure 4. Such internal models are referred to as *forward models* in control systems theory. We assume that the motor latency *T*_mot_ largely represents delays in transmitting the control signal from the brain to the eye. We assume that the virtual plant is located in the brain close to where the neural control signal is generated, and thus has access to this signal with negligible delay. Accordingly, the output of the virtual plant is *predicted future accommodation*, i.e. the value that ocular accommodation will have at a time *T*_mot_ in advance of the present. We write this predicted future accommodation as *â*(*t*+*T*_mot_): the predicted accommodation at a time *T*_mot_ in the future, where the circumflex indicates that this is an *estimate* of the future accommodation. Since the accommodation up to a time *T*_mot_ into the future is controlled by neural signals already sent by the brain, this estimate can in principle be perfect. It should be affected only by noise, and by any inaccuracies in the virtual plant as a model of the ocular plant. In the model we present here, neither of these apply and so the prediction of future accommodation is indeed perfect.

Predicting stimulus demand is more challenging, since in general this reflects the motion of objects in the outside world. Nevertheless, several studies (Campbell & Westheimer, 1960; Charman & Heron, 2000; Krishnan et al., 1973; Phillips et al., 1972; Stark et al., 1965) have suggested that the accommodation system, like vergence and other motor systems (Erkelens, 2011; Rashbass & Westheimer, 1961), may be capable of predicting sufficiently regular input. For example, if the demand is a square wave, jumping between two values with a constant period, accommodation develops a very short latency or even changes in anticipation (Krishnan et al., 1973). How or whether this prediction is achieved is beyond the scope of this paper; it may be performed by the cerebellum (Ohtsuka & Sawa, 1997; Popa & Ebner, 2019) or it may not actually occur (Águila-Carrasco & Marín-Franch, 2021; Otero et al., 2019). The different possibilities can be modelled with the *Demand Predictor* block (Figure 4). This takes as its input what demand is estimated to have been at time *T*_sens_ in the past, 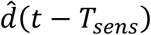, and gives as output what it estimates demand will be at time *T*_mot_ in the future, 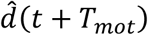. That is, it extrapolates its input into the future by a time corresponding to the entire sensorimotor latency, *T*_lat_*=T*_mot_+*T*_sens_. In this paper, our model Demand Predictor block will simply pass its input on unchanged, effectively assuming that the demand will stay at its current value. This is probably a reasonable assumption, since in many natural viewing situations, accommodative demand probably often changes rather little over the timescale of *T*_lat_. A future model could incorporate a more elaborate form of prediction, e.g. taking account of stimulus periodicity, but that is beyond the scope of this paper.

Having introduced the key elements of the predictive model – the virtual plant and the demand predictor – we now discuss how it works. To help with this, we have annotated the signals in Figure 4 and marked some reference points with red letters. Let’s start at A with the output of the virtual plant. As we saw above, this represents the brain’s prediction of what ocular accommodation will be at time *T*_mot_ in the future: 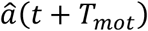. Our model brain uses this predicted future accommodation in two ways. First (B), the model brain delays this predicted-accommodation signal by the total sensorimotor latency to obtain 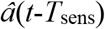, an estimate of what the ocular accommodation was at a time *T*_sens_ in the past. Thus, the predictive model actually uses an internal estimate of *past* accommodation as well as of future accommodation. The point of doing this is to match the latency of the defocus signal. The input to the whole system is accommodative demand, *d*(*t*) (label D). In the eye (label E), the ocular accommodation *a*(*t*) is optically subtracted from *d*(*t*) to yield the error signal *e*(t), the optical defocus at time *t*. Ideally, this is what the accommodation control should be based on, but due to the sensory latency *T*_sens_, the brain only has access to the delayed signal, *e*(*t-T*_sens_), representing the defocus at a time *T*_sens_ in the past. At the signal combination labelled C, the brain adds its estimate of past accommodation, 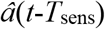, back onto this delayed defocus signal *e*(*t-T*_sens_), in order to obtain an estimate of what the demand was at a time *T*_sens_ in the past: 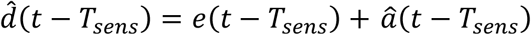. This demand signal is fed into the Demand Predictor block, which uses it to make a guess at what the demand will be at a time *T*_mot_ in the future: 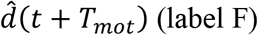.

Now, the brain makes its second use of predicted future ocular accommodation, this time without applying any delay. At the signal combination labelled G, it subtracts the predicted accommodation 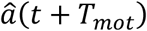 from the predicted demand 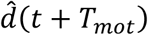 to obtain the predicted future defocus error: 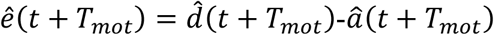. This predicted future defocus is what is fed into the yellow Controller block and used to compute the neural control signal driving accommodation. It is this use of predicted future defocus which makes this a predictive model, as compared to the model shown in Figure 3.

As noted above, a constant bias is added on to the output of the controller, which accounts for the non-zero resting focus. We call the result *m*(*t*) (label H). This is the actual motor signal sent to the ocular plant, with a latency *T*_mot_, which results in the ocular accommodation *a*(*t*) (label I). An efference copy of the same motor signal is also sent to be the input of the virtual plant. The output of the virtual plant is, of course, the predicted future accommodation that we began with (A), so we have now followed the signals around the whole of the inner and outer loops.

In summary, then, although the input to the accommodative control system as a whole is the sensed *current* defocus (Figure 2), in a predictive model the input to the accommodative controller itself is the predicted *future* defocus. With this modification, PID-type controllers can now work well and avoid the instabilities associated with an out-of-date error signal.

#### Simplified representation of the predictive control system

It is useful to note that, if the virtual plant is a perfect simulation of the physical plant, the predictive control system shown in Figure 4 is mathematically equivalent to the much simpler form shown in Figure 5. This form can appear confusing, because it shows accommodation being subtracted from the stimulus demand *after* the sensory latency (even though some of the sensory delay represents the optic nerve and cortical processing) and *before* the motor latency (even though that represents processes before accommodation). The reader is invited to trace the signals around Figure 4 and Figure 5, and verify that provided 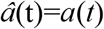, the same inputs are fed into the same blocks and so the results must be the same. Figure 5 provides a visual picture of what is being achieved by the predictive control: it effectively shifts the latencies outside the control loop. This diagram holds whatever the demand predictor does. If the demand predictor were able to predict future demand perfectly, it would cancel out the latencies and the system would behave as if there were no latencies. But even if the demand predictor merely assumes demand stays constant, as in our model, it still makes the control immune to the destabilising effect of latencies. The effect of latencies is now only to delay the response. The response to any stimulus is exactly the same as for a system with perfect prediction of demand, just occurring later in time (see Appendix and Table 3). Thus although predicting the sensory input enables a more rapid response, predicting one’s own motor response suffices to ensure stability.

**Figure 5.**
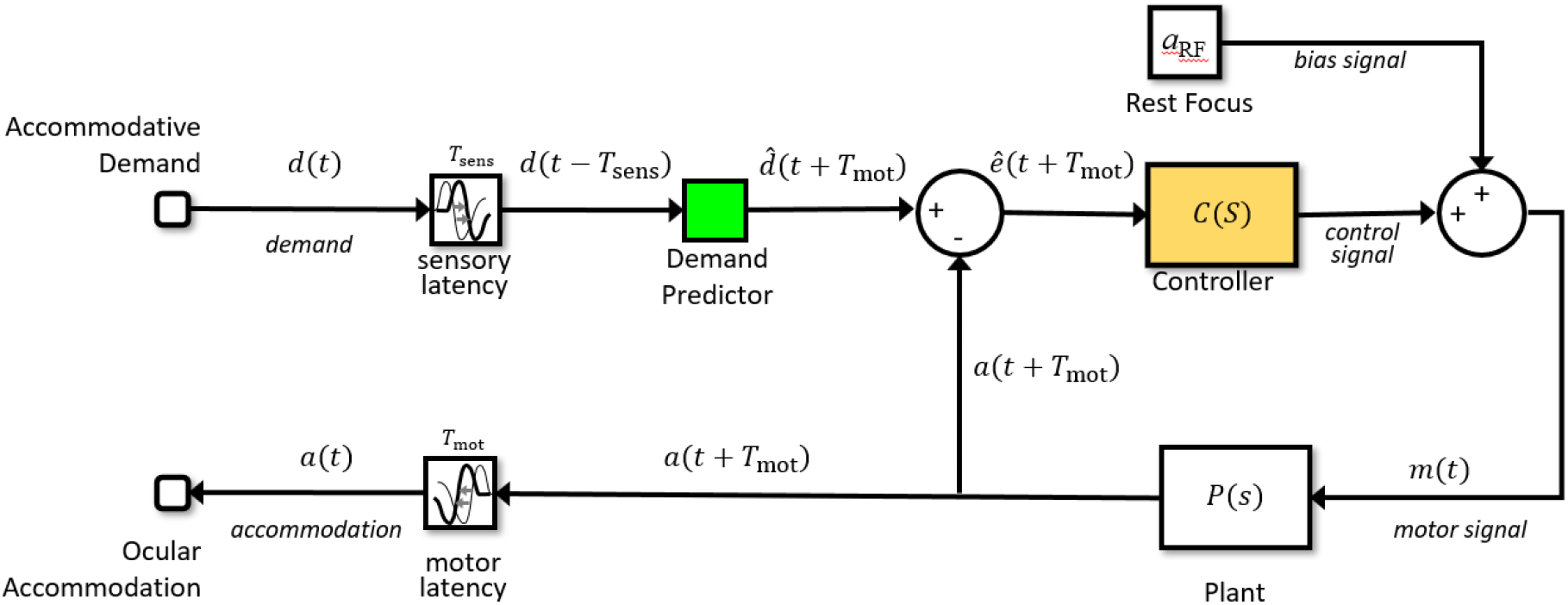
*Simplified version of the model shown in Figure 4. This “non-causal” model structure is not physiological and cannot be mapped onto “brain” and “eye” like the predictive physiological model in Figure 4. For example, here the single block labelled “Plant” is used to represent both the physical plant in the eye and the virtual plant modelled in the brain. However, as shown by the annotated signals, it is mathematically equivalent to the physiological model in Figure 4*, provided *that the Virtual Plant block is a perfect simulation of the Ocular Plant*.

**Table 3.**
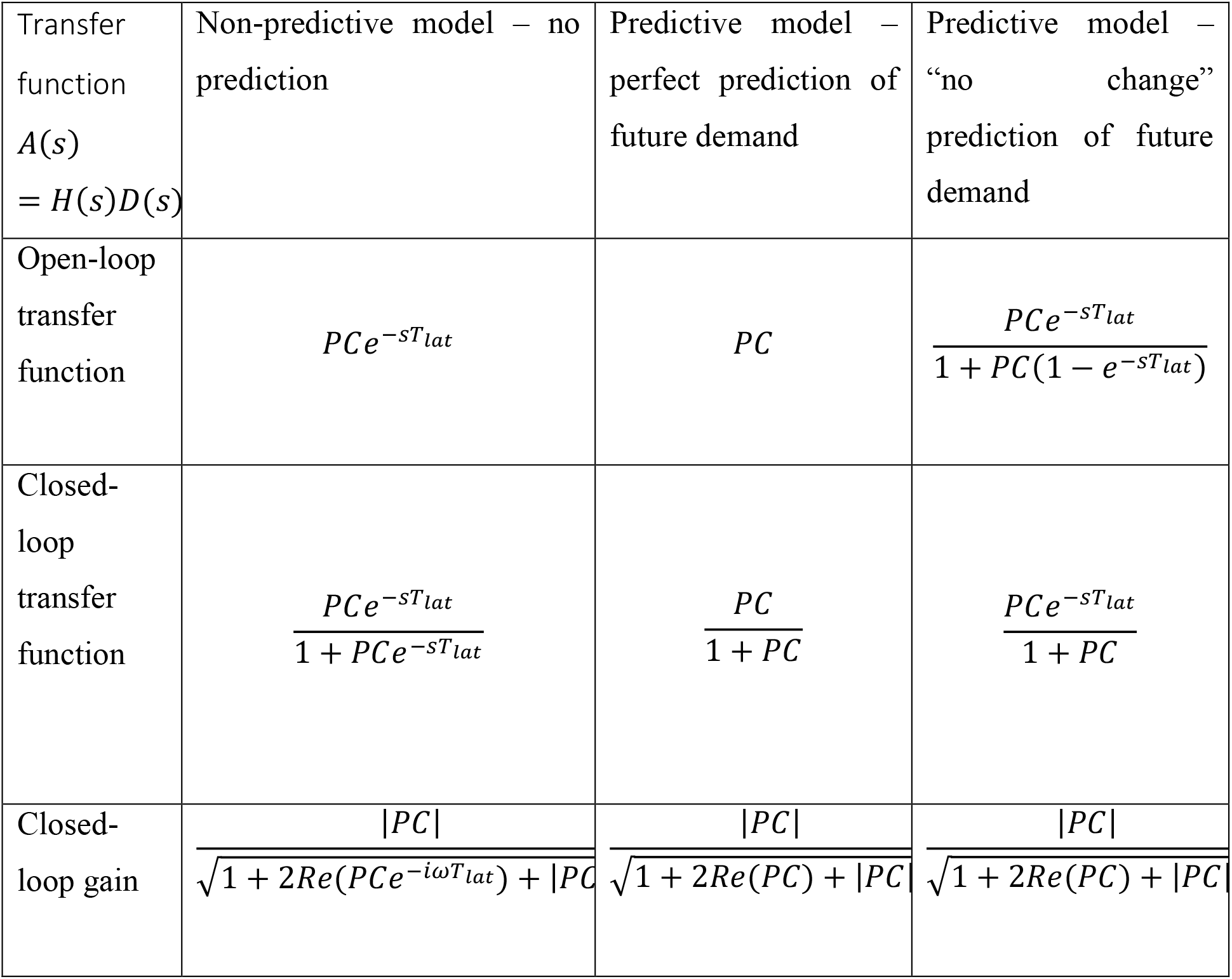
Open- and closed-loop transfer functions H(s) for different control systems; see Appendix for derivation. The transfer function relates accommodation to the demand via A(s) = H(s) D(s), where A(s) is the Laplace transform of accommodation relative to rest focus, a(t)-a_RF_, and D(s) is the Laplace transform of demand relative to rest focus, d(t)-a_RF_. P(s) the transfer function of the ocular plant, and C(s) is the transfer function of the neural control (block marked Controller in Figure 3, Figure 4, Figure 5). T_lat_ is the total sensorimotor latency from a change in demand to the accommodative response.

### A specific model of accommodative control

So far we have deliberately kept the discussion very general, without committing to a particular choice of transfer function for either the ocular plant or the Controller block which converts defocus into a neural signal to the plant. In this section, we develop and justify a more specific model of accommodative control. We discuss plausible assumptions and constraints on both the forms of these transfer functions, and their particular parameters.

#### Ocular plant

The Ocular Plant block in Figure 2–Figure 5 converts the motor neural control signal *m* into accommodation *a*. Physiologically, this block corresponds to the following components. The ocular lens is held in an elastic capsule between the anterior and posterior chambers of the eye. It is tethered along its equator by elastic suspensory ligaments or zonules. The axial zonules pass from the lens equator to the inner margin of the ciliary muscle, while the posterior zonules pass from the ciliary muscle back to the choroid at the ora serrata, the junction between the choroid and the ciliary body. The lens is flattened by the elastic tension under which it is held by the zonules, and becomes more spherical – and so more optically powerful – when its extension is reduced by the constriction of the ciliary muscle. Figure 6A shows a diagram of this arrangement. Figure 6B shows a simplified biomechanical model (Beers & van der Heijde, 1994, 1996; Schor & Bharadwaj, 2005; Wang & Pierscionek, 2019). The zonules, choroid and ciliary attachment are represented as springs. The lens is represented by a Voigt model, in which a spring is in parallel with a dashpot or damper. The springs are modelled according to Hooke’s law, i.e. they exert a force proportional to their extension. The dashpot exerts a force proportional to the rate of change of its extension, modelling the viscosity of the lens and capsule. The whole system is subject to the force *f* exerted by the ciliary muscle, which is set by the neural signal sent by the accommodative control system. We assume that the optical power of the lens is proportional to the extension of the spring/dashpot modeling the lens.

**Figure 6.**
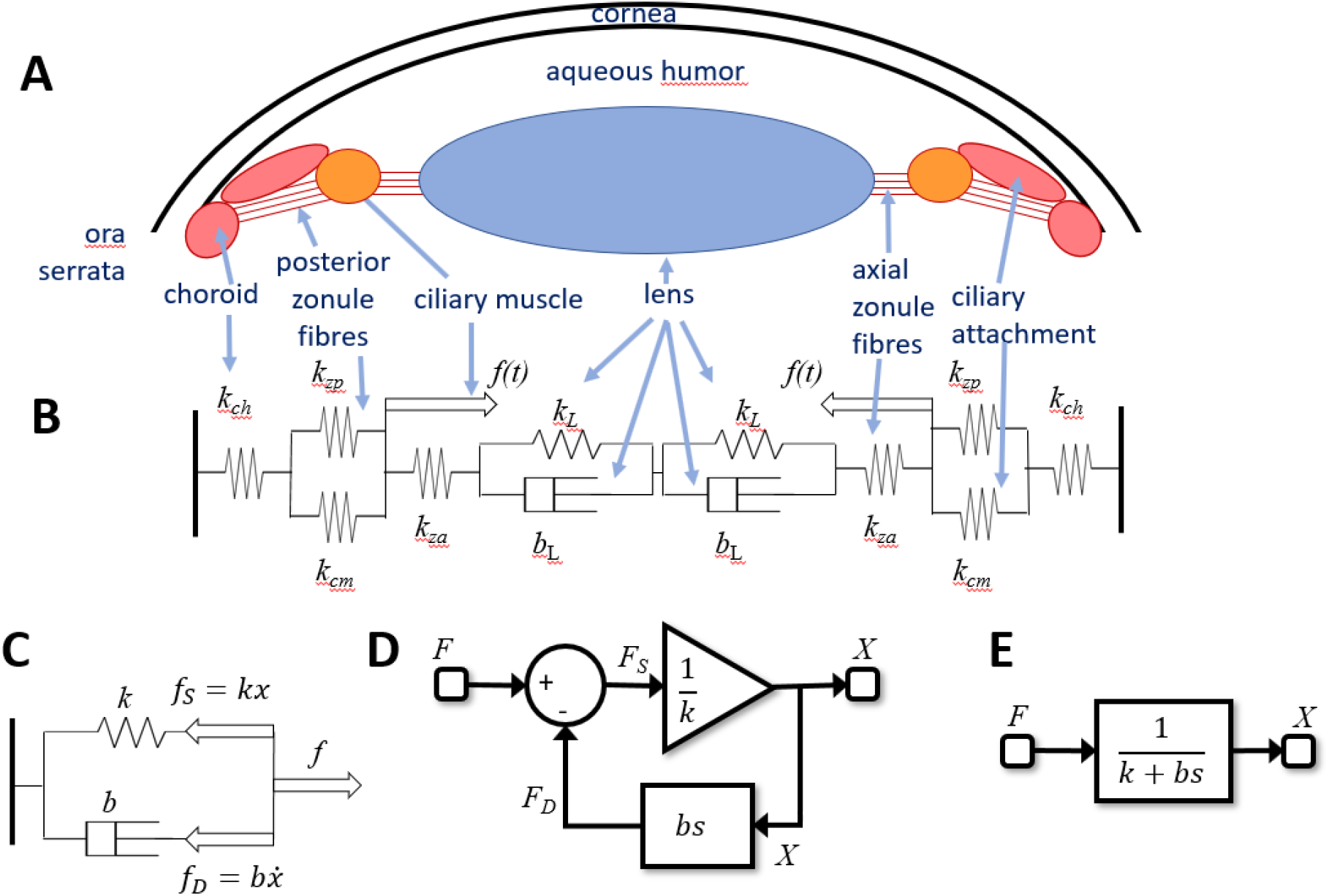
A: Diagram of the anatomical structures relevant to accommodation. B: Representation as a biomechanical model, consisting of a set of elastic springs (spring constant k) and dashpots (viscosity b). The posterior zonule fibres and ciliary attachment are assumed to be in parallel, so their extensions are equal. C: Minimal model which is mathematically equivalent to the full model shown in B. The parameters k and b are functions of the original parameters. The thick arrows mark forces. As well as the ciliary muscle force, we now have the force in the spring, f_S_, and in the dashpot, f_D_.. D: Control theory block diagram equivalent to the simple model in C. For example, at the summation block, we have the force balance F_S_=F-F_D_; at the gain block, we have X=F_S_/k; in the feedback loop we have F_D_=bsX. E: Single transfer function equivalent to the block diagram shown in D. This is a leaky integrator, with time-constant τ_plant_=b/k.

Since by Newton’s laws the forces must sum to zero at every point, the system shown in Figure 6B represents a set of simultaneous equations; for example at the junction between the axial zonules and the lens, we have

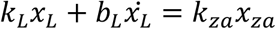

where *x*_L_, *x*_za_ are the extensions of the lens and of the axial zonules respectively, *k* their spring constants and *b*_L_ the viscosity of the lens. Using the constraint that the sum of all the extensions must be constant, we can go through and solve the simultaneous equations for the lens extension *x*_L_. If we do so, the result is the same as for the simplified system shown in Figure 6C, with a dashpot and a single spring, now representing the combined elasticity of all the component elements. The value of the full model is that the elasticity of the different tissues can be measured independently. This is important if one wants to model age-dependence (Schor & Bharadwaj, 2005), since these vary differently with age, but the collapsed model is obviously much simpler to work with.

In control theory, a spring can be viewed as an LTI element converting an input, force, into an output, extension. The transfer function mapping force to extension is thus simply the inverse of its spring constant *k*, i.e. its compliance. A dashpot is similar, but since the force is proportional to the rate of change of extension, the transfer function mapping extension to force is *bs*, where *b* is the viscosity and *s* represents differentiation (see primer above). In this way, the simple biomechanical model shown in Figure 6C can be represented by the block diagram in Figure 6D, or even more succinctly by the transfer function in Figure 6E. This is the transfer function of a first-order low-pass temporal filter with time-constant τ_plant_=*b/k*, also known as a leaky integrator. This, then, is the function mapping ciliary muscle force to lens extension.

We now make two further simplifying assumptions : (1) that the brain is able to command ciliary muscle force directly, so that the motor signal sent to the plant from the brain can be regarded as proportional to ciliary muscle force, and (2) that optical power is proportional to lens extension. With these assumptions, then, the entire ocular plant block mapping neural signal to accommodation can be regarded at least roughly as a leaky integrator (Beers & van der Heijde, 1994, 1996; Ejiri et al., 1969; Wang & Pierscionek, 2019). We therefore model the transfer function of the plant as

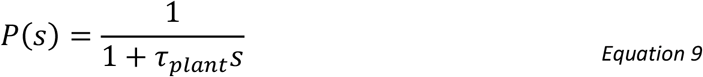

where empirically τ_plant_ is around 0.156s for young eyes (Schor & Bharadwaj, 2006). In this paper, we will take this value as a given. As noted above, we can assume without loss of generality that the steady-state gain is 1.

#### Controller

We now come to a key decision: the choice of transfer function for the Controller, *C(s)*. As noted above, in industrial control systems, controllers typically have proportional, integral and derivative (PID) terms, with transfer functions which scale as constant, 1/s or s respectively.

We can rule out pure proportional control, since with *P*(s) as given in Equation 9, making *C(s)* constant means that the system tracks rapid sinusoidal oscillations far better than human accommodation. For example, *C(s)*=5 results in a realistic steady-state gain of 83% (Equation 7), but the gain remains >50% out to frequencies as high as 8Hz, far higher than observed (see Figure 7 below). Derivative terms do not affect steady-state error, but improve stability and avoid overshoot. They also enable rapid response to rapid changes. However, they can be problematic in the presence of noise. Previous work by Schor and Bharadwaj (Bharadwaj & Schor, 2006; Schor & Bharadwaj, 2004, 2006) suggests that the accommodative system has a distinct “pulse” mechanism for responding to sudden large changes in accommodation such as occur when we change from looking at a distant to a near object, which cannot be modelled by an LTI system and which are beyond the scope of this paper. Furthermore, many of the benefits of derivative control are already achieved by our use of a forward model to predict future demand. We therefore do not include a derivative term. This leaves us with the integral term. A pure integral controller has a transfer function proportional to 1/s, and thus infinite gain at s=0. This is desirable since it eliminates steady-state error, but it also means that errors can accumuluate; also as noted, the human accommodation does not seem to completely eliminate steady-state error. We can account for this by modelling the controller as a leaky integrator, following Krishnan and Stark (1975):

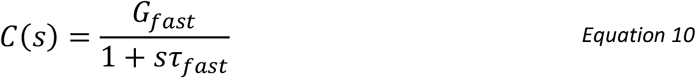

where G_fast_ is the steady-state gain and τ_fast_ the time-constant. The subscript “fast” is to distinguish this from a slow integrator which we shall introduce below. A leaky integrator acts like a pure integral controller over short timescales (*s*τ>>1), and like a pure proportional controller over long timescales (*s*τ>>1), thus combining aspects of both. We noted above that accommodative lead/lag suggests the steady-state gain must be in the range 4-9. We somewhat arbitrarily chose *G*_fast_=8.

**Figure 7.**
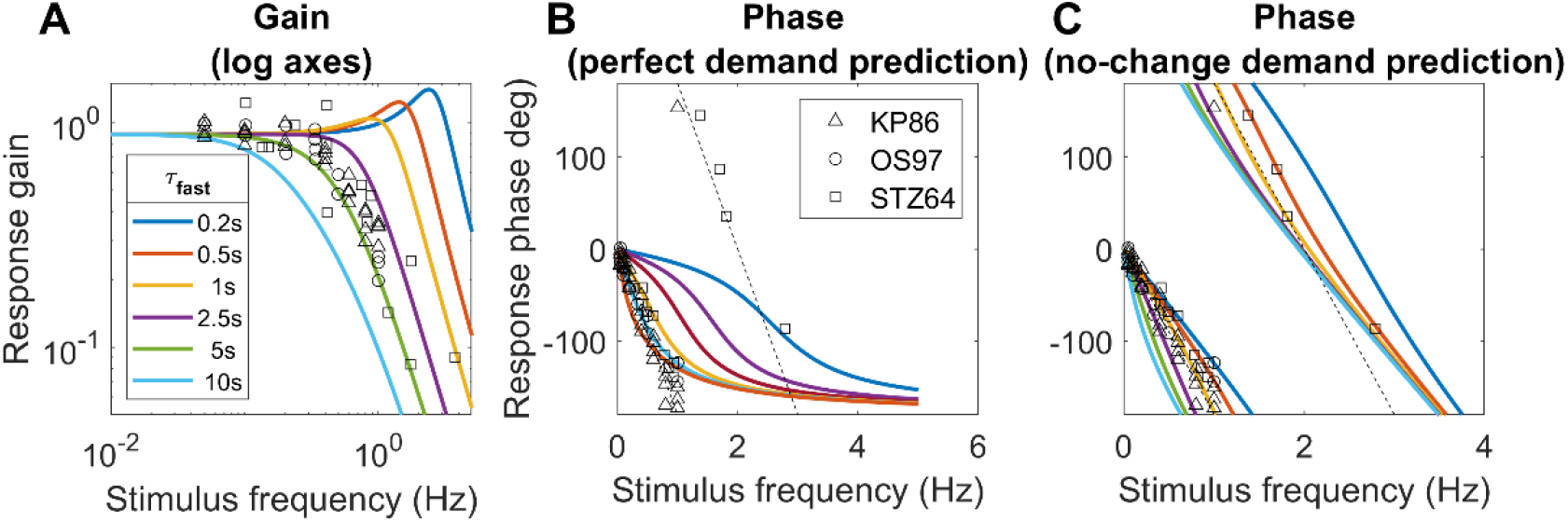
Constraints on the time-constant of the fast integrator. Coloured lines show the gain and phase predicted for a predictive model with leaky-integral control, Table 3, with P(s) given by Equation 9 with τ_plant_=0.156s, and C(s) given by Equation 10 with G_fast_=8 and different choices of τ_fast_. The phase is shown (B) for a model capable of predicting demand perfectly, or (C) for the “no-change” model which simply assumes demand will continue at its instantaneous value; both of these have the gain shown in A. Symbols show empirical results from Kruger and Pola (1986), Ohtsuka and Sawa (1997), and Stark et al (1965). The dashed line in the phase plots corresponds to a constant latency of 0.5s, close to what is observed. Code to generate this figure is in Fig_TimeConstraints.m.

#### Gain for sinusoidal input: sub-critical damping

With both the plant and the controller being leaky integrators, and with a predictive control system, the closed-loop gain is that of a damped harmonic oscillator (Equation 19, Appendix). The behavior of this system can be summarized by its natural frequency and damping coefficient ζ, both of which depend on the parameters G_fast_, τ_fast_, τ_plant_ (Equation 20). If the damping coefficient ζ is too low, the maximum gain is observed for a non-zero resonance frequency, and can even exceed 1. This does not agree with empirical observations of accommodative response to sinewaves, which is low-pass (Charman & Heron, 2000; Kruger & Pola, 1986; Ohtsuka & Sawa, 1997; Stark et al., 1965); Figure 7A. This indicates that ζ is at least 1/√2, not far below critical damping (ζ=1) (Labhishetty & Bobier, 2017). Saccades have a damping coefficient of around 0.7 (Bahill et al., 1975); systems with this value have minimum settling time, i.e. they reach and remain within 5% of their final value most rapidly. We show in the Appendix that obtaining ζ~1/√2 for a system with G_fast_>>1 requires the time-constant of the fast controller to be

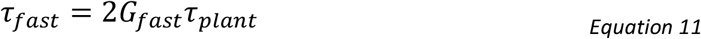

Thus, with τ_plant_=0.156s and G_fast_=8, τ_fast_ must be at least 2.5s.

#### Phase for sinusoidal input: further evidence for predictive control

Empirically, up to ~1Hz the phase delay of accommodation is very close to a linear function of frequency, indicating a constant latency *T*_delay_ : *ϕ* = 2*πfT*_delay_ (Charman & Heron, 2000; Heron et al., 1999; Kruger & Pola, 1986; Ohtsuka & Sawa, 1997; Wildt et al., 1974). The slope usually corresponds to a delay of ~0.5s (dashed lines in Figure 7BC), though there is considerable variability between studies. Because 0.5s is close to the sensorimotor latency inferred from the response to step changes, it is often therefore assumed that this phase slope must represent the sensorimotor latency. However, this is not necessarily the case. First, the damped second-order system formed by the ocular plant and the neural control imposes delays in addition to the sensorimotor latencies. Second, if the brain predicts demand perfectly – at least theoretically possible for a regular stimulus like a sinewave – then its phase delay becomes independent of the sensorimotor latency (see Appendix).

#### The time-constant of the fast integrator

Thus, together the gain and phase response of accommodation to sinusoidal oscillations in demand place quite tight constraints on the time-constant of the fast integrator, τ_fast_, given that the time-constant of the plant is a biomechanical given, and the gain of the fast integrator is already quite tightly constrained by the observed lead/lag following a change in demand. Figure 7 illustrates this by comparing the theoretical gain and phase with different values of τ_fast_ with empirical results from various subjects and studies. As noted, we can rule out τ_fast_ < 2.5s because the gain is then too high at high frequencies. The gain data is probably best described by τ_fast_ = 5s (green lines in Figure 7A), but this does not account for the phase data. τ_fast_ = 5s in the perfect-prediction model gives phases which match empirical data up to around 0.5Hz, but at higher frequencies, empirical phase continues to increase roughly linearly, implying a constant delay, whereas phase for the perfect prediction model asymptotes at 180° (Figure 7B). Thus, we probably have to reject the perfect-prediction model (not surprising given its idealised nature). The no-change prediction model is qualitatively in much better agreement with the phase data, but then τ_fast_ = 5s predicts larger phases than are observed (Figure 7C). The purple line shows the curve with minimum settling time, τ_fast_=2.5s which yields ζ~1/√2. This is in reasonable agreement with both gain and phase data, assuming simple no-change demand prediction, and we therefore adopt this value in the rest of the paper.

#### Adaptation and dual control

Another distinctive feature of accommodation is that it adapts after prolonged exposure to the same demand. This can be revealed by using pinholes to place the system in open-loop mode. As we have seen, in this situation, accommodation returns to the resting focus. After short periods of stimulation, this happens rapidly, in a few seconds. However, after long periods of exposure to a particular demand, the return happens over a much longer time period, sometimes several minutes. This cannot be accounted for with the leaky-integral control proposed so far. However, it can be explained by positing a dual control system in which a fast, or phasic, neural integrator controls changes in response amplitude and a slow, or tonic, neural integrator maintains the response amplitude (Khosroyani & Hung, 2002; Schor, 1979a; Schor et al., 1986; Sun & Stark, 1990).

The fast integrator is the one we have considered so far, which responds to error signals computed from negative feedback. The slow integrator responds to the activity of the fast neural integrator, and not directly to the error signal. As the name implies, the slow integrator has a long time constant, which means that it has little effect on the response to rapid changes in demand, so our previous discussion of the responses to sinusoids is not invalidated by its addition. With this arrangement, the transfer function of the Controller becomes

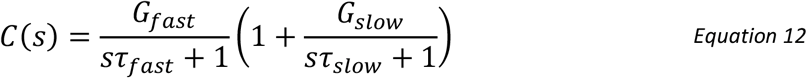

The steady-state open-loop gain of the system is therefore

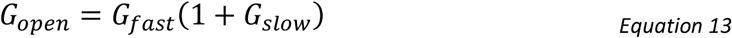

Figure 8 shows the non-predictive control system of Figure 3 after the addition of this second, slow integrator, since it is easier to appreciate its operation in a non-predictive system. Suppose the system starts from rest, with demand and accommodation both equal to the rest focus, so that the defocus error and the outputs of the fast and slow integrators are both zero and the neural signal sent to the plant is simply the bias signal, maintaining it at the rest focus. Suppose the demand then makes a step change to a nearer value, d_0_. This in turn makes the defocus error non-zero, which begins to charge up the fast integrator. The output of the fast integrator increases the neural control signal above the bias value, altering accommodation so as to reduce the error. It also begins to charge up the slow integrator. Thus, over short timescales, the neural signal controlling accommodation is set mainly by the output of the fast integrator. However, over long timescales, the slow integrator takes over. The ratio of the slow to fast steady-state contributions is equal to the gain of the slow integrator (Schor, 1979b; Schor et al., 1986); for example, with our value G_slow_=5, steady-state accommodation is 83% due to the slow integrator and 17% due to the fast integrator.

**Figure 8.**
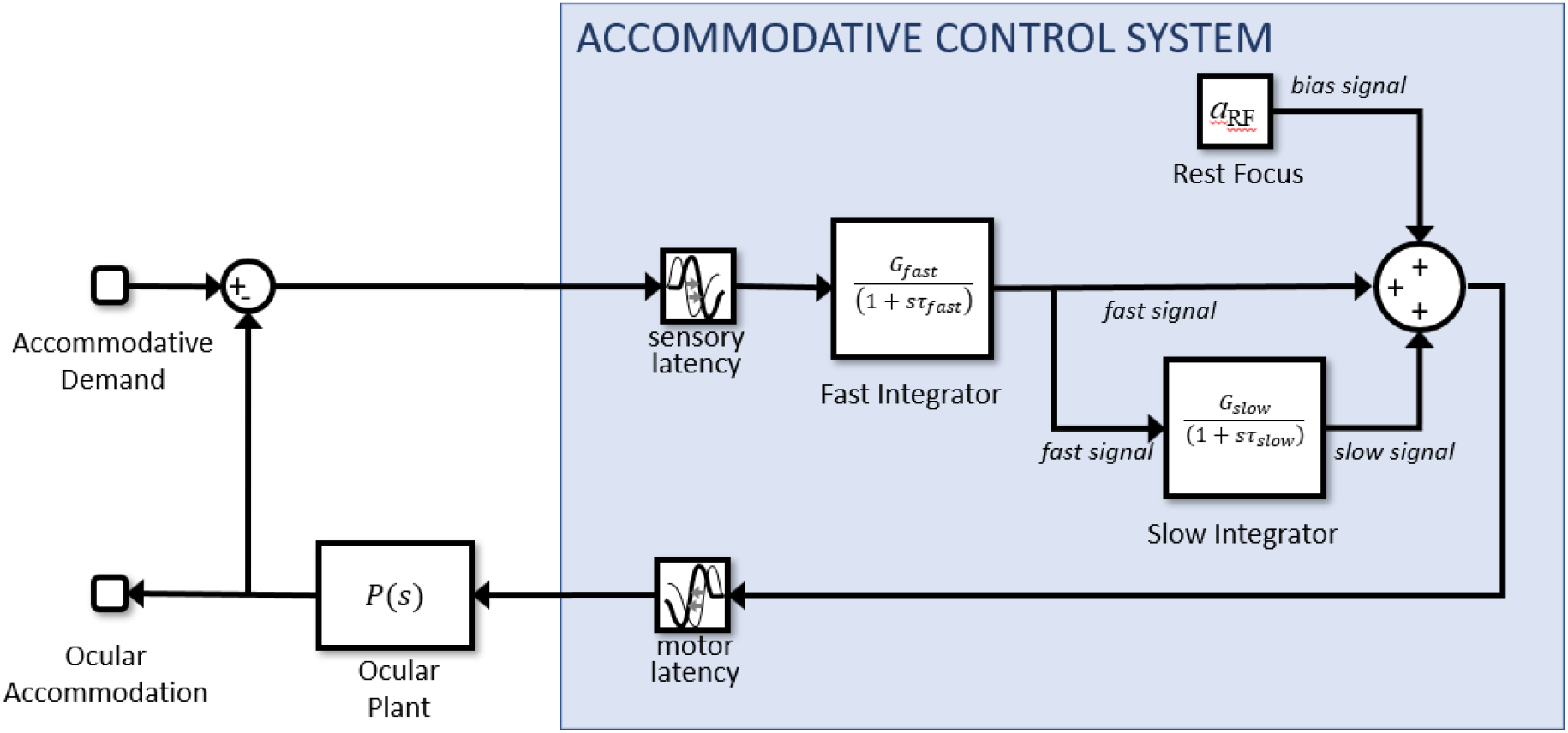
Non-predictive model incorporating dual (fast+slow) control. The slow integrator can be added to predictive models in the same way, but its effect is then much more complicated.

Now suppose that pinholes are applied, making the defocus error zero regardless of accommodation. In this non-predictive model, after a delay corresponding to the sensory latency, the signal entering the fast integrator instantaneously drops to zero, and the fast integrator begins to discharge. As the fast integrator discharges, accommodation drops rapidly, with a decay time corresponding to τ_fast_. When the signal from the fast integrator has dropped far enough, the slow integrator begins to discharge as well, resulting in a second, slower decay of accommodation, with a time constant corresponding to τ_slow_. Thus, after a long period of exposure, there is an initial rapid drop as the proportion of accommodation due to the fast integrator, initially 1/(G_slow_+1), decays rapidly, but then a much longer decay as the dominant component due to the slow integrator decays slowly.

The slow integrator also increases the overall steady-state gain and thus reduces the steady-state error. Using Equation 13 and Equation 7, the steady-state accommodative response is

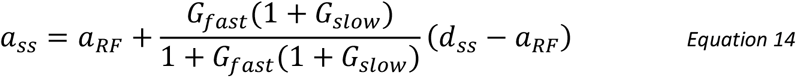

where with *G*_fast_=8, *G*_slow_=5 the gain term is 0.98, compared to 0.89 with only the fast integrator. Thus, following a step-change in demand, the model response rises rapidly to around 90% of the demand, and then over the next tens of second rises more slowly to approach the demand exactly. Thus, the gain of the slow integrator cannot be made too large (say, much larger than 5) without eliminating the ability of the model to account for accommodative lead and lag.

With predictive control, there is an additional subtlety which also places an upper bound on *G*_slow_. In such systems, the fast integrator is driven not by retinal defocus directly, but by the estimated future defocus (Figure 4). This does *not* immediately drop to zero when pinholes are applied. When the system is made open-loop by setting *d(t)*=*a(t)*, the input to the fast integrator becomes *a*(*t-T*_sens_)-*a*(*t*+*T*_mot_) for the no-change prediction model. This becomes zero once accommodation has stabilized, but is finite while it decays. When the gain of the slow integrator is sufficiently large, this small error input is enough to keep the slow integrator high. This in turn keeps accommodation high and thus sustains the error signal. Accommodation creeps slowly down to the rest focus with a time-constant which, counter-intuitively, can be much longer than any of the three time-constants of the system: τ_plant_, τ_fast_, τ_slow_. This effect is independent of exposure duration, so cannot account for the adaptation which the slow integrator was introduced to explain. To avoid this effect and obtain a clear difference between short and long exposure durations, we have found that *G*_slow_ needs to be less than around 10. Here, we have set *G*_slow_=5.

#### Microfluctuations and noise

A distinctive property of accommodative response is the relatively large fluctuations to which it is subject in both open and closed loop. The power spectrum of open-loop accommodation is roughly a straight line on log-log axes (Campbell et al., 1959b; Campbell & Westheimer, 1960; Stark et al., 1965), i.e. a power-law spectrum, P=1/*f*^α^. We model this by injecting white noise onto the defocus signal prior to input to the neural controllers (Figure 10). White noise has a flat power spectrum, but integration by the two integrators within the system (the neural controller and the plant) converts it to a power-law spectrum, with an approximately Brownian (1/*f*^2^) spectrum.

Noise has often been omitted from models of accommodative control, presumably with the rationale that once the correct noise-free response has been obtained, noise can always be added later to simulate microfluctuations. However, this approach is unwise, because noise in fact adds important constraints to the system. This is especially true with a predictive control system, which can easily end up amplifying noise in the open-loop condition. Referring to Figure 4, we see that a predictive control system contains not one but two feedback loops: one via the eyes, and one internal to the brain, incorporating the virtual plant. Operating in open-loop mode cuts the outer feedback loop, but leaves the internal feedback loop intact. Depending on the coefficients, internal noise can easily resonate within this loop, creating a situation where the power spectrum of open-loop accommodation has sharp peaks which do not occur in closed-loop mode, since the outer feedback loop suppresses them in its effort to keep the error zero. This is not observed empirically. The power of low frequencies does increase in open-loop mode (Charman & Heron, 2015; Gray et al., 1993b), since without an error signal accommodation performs a random walk around the rest focus, whereas it is kept close to the demand in closed-loop mode. But we do not see an increase in the power of particular high-frequencies, as would occur if internal noise were resonating within the internal feedback loop. Fortunately, we find that the values we have already derived are consistent with these data. A more underdamped system – say *G*_fast_=15, τ_fast_=2s, which puts the damping coefficient ζ at 0.5 – does show unrealistic high-frequency resonances within the forward model feedback loop, but our sub-critically-damped parameters *G*_fast_=8, τ_fast_=2.5s, ζ=0.7 already suppress the open-loop resonance.

#### Explaining the closed-loop resonance seen for high frequencies at low amplitudes

In fact, several workers have found evidence for a resonance in closed-loop but *not* open-loop mode. The first evidence comes from microfluctuations during steady fixation. Several workers have found that the power-spectrum of closed-loop accommodation has a peak at around 2Hz (Figure 9A). It is not always present, but when found is always *more* prominent in closed-loop than open-loop accommodation. Although the location of this peak varies with heartrate, suggesting the pulse as a possible source interacting with blood volume of the ciliary body (Collins et al., 1995; Winn et al., 1990), the fact that it is higher in closed-loop conditions suggests that the source must be amplified by a neural resonance within the outer feedback loop.

**Figure 9.**
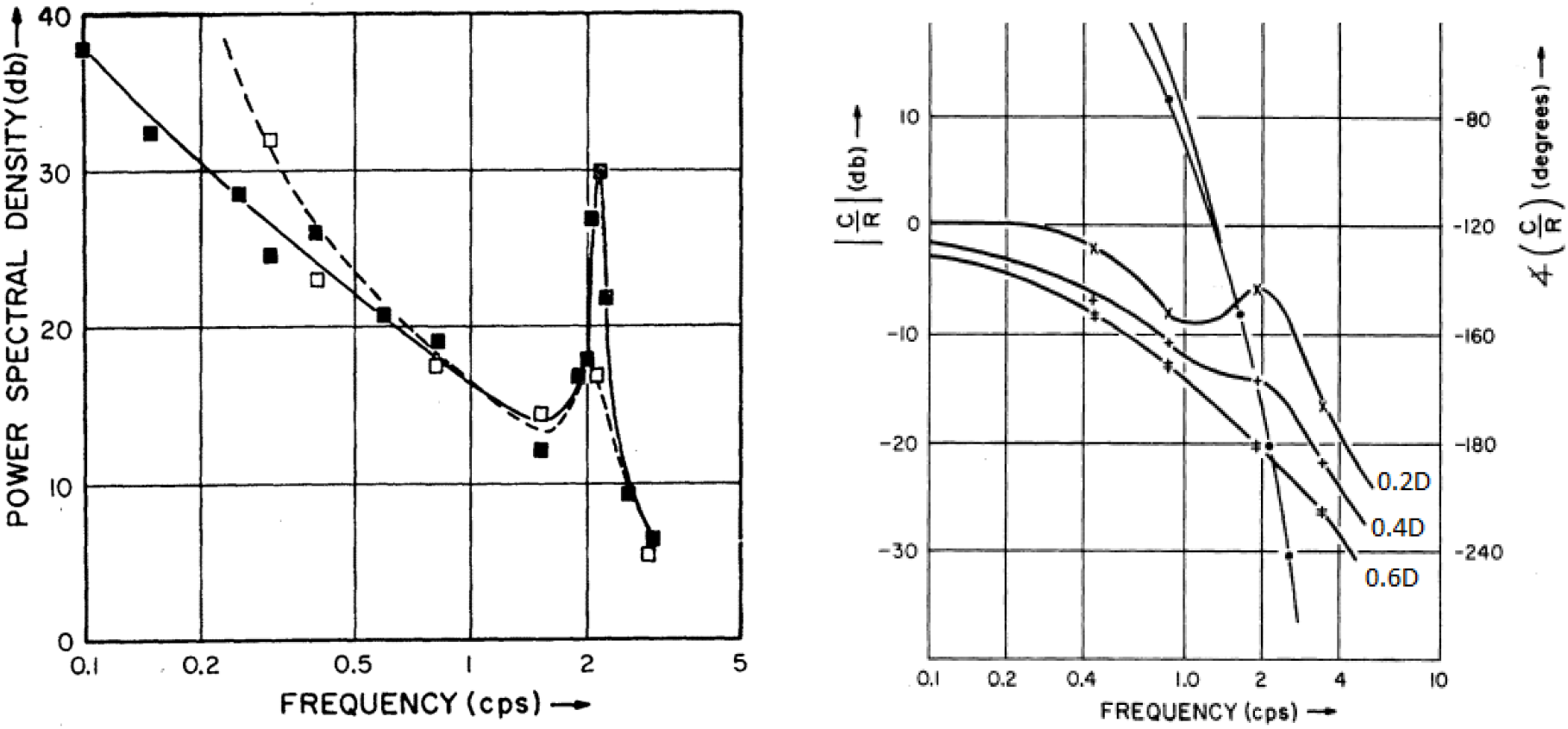
Evidence for a resonance at around 2Hz in accommodative control. (A) Figure 5 from Stark et al 1965 (Stark et al., 1965), replotting empirical results from Campbell et al 1959 (Campbell et al., 1959b), showing the power spectrum of accommodation under closed-loop (solid) and open-loop (pinhole, dashed) conditions. (B) Empirical results from Figure 4 of Stark et al 1965, showing gain for sinusoidal oscillations of three different amplitudes (0.2D, 0.4D, 0.6D). Gain is expressed in decibels (left axis): 0dB corresponds to an amplitude gain of 1, −10dB to 0.32, −20dB to 0.1, −30 to 0.03.

Furthermore, the same resonance is assumed to be responsible for another puzzling observation, relating to gain with sinusoidal stimuli. In our discussion around Figure 7, we emphasized the lowpass nature of the gain response. This is true at high amplitudes, but for low-amplitude oscillations in demand, the curves become non-monotonic, with an increase in gain at around 2Hz (Figure 9B). Ockham’s Razor suggests this reflects the same closed-loop resonance causing the ~2Hz peak in microfluctuations. However, the dependence on amplitude indicates that this resonance must be caused by a nonlinear mechanism, since for a linear system gain is independent of stimulus amplitude.

Resonances observed in closed-but not open-loop mode immediately suggest a control system lacking the predictor we have argued for so far. Non-predictive control is prone to closed-loop instabilities in systems with latencies, like accommodation. This occurs in the outer feedback loop via the eye, when the accommodation change designed to null out defocus arrives out of phase due to the latency and ends up enhancing the defocus which cause it. Predictive control avoids these closed-loop instabilities, but if the prediction is imperfect, it can be vulnerable to open-loop resonances due a similar effect occurring via the internal feedback loop driven by the efference copy. (For a mathematical justification of these statements, see the Appendix, specifically the discussion around Equation 15, Equation 16 and Equation 18.)

Thus to explain both the power spectrum of microfluctuations, and the non-linear resonance in the response to sinusoidal demand, we postulate an additional signal controlling accommodation. This is proportional to small amplitudes of the current defocus, not the estimated future defocus, and is thus not predictive. (This signal is, however, included within the efference copy used to estimate future defocus within the predictive control system, as shown in Figure 10). Because this signal is non-predictive, it is prone to closed-loop instabilities. But for the same reason, it avoids open-loop resonances which can occur within a predictive system.

**Figure 10.**
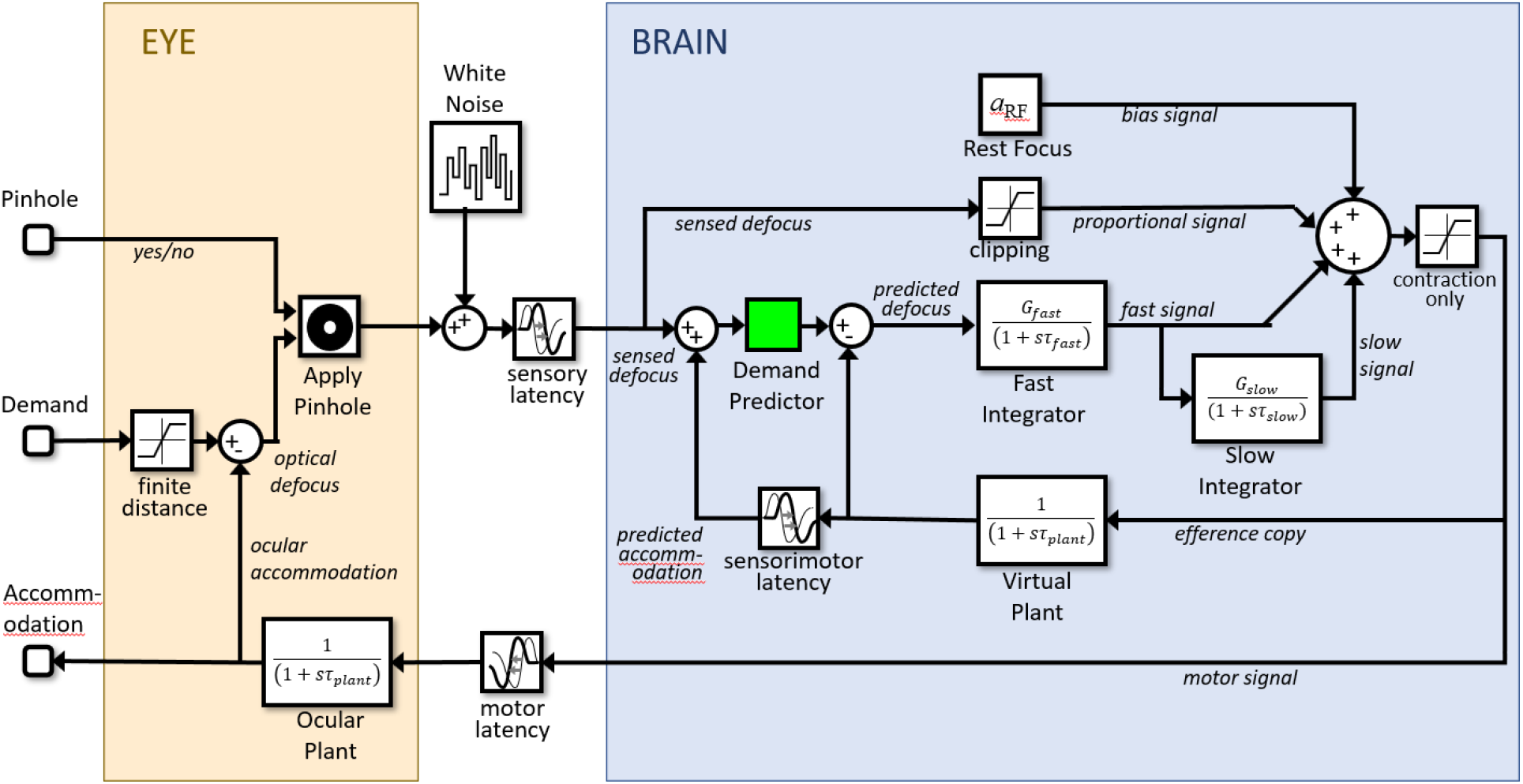
Block diagram of our final model (Simulink file AccommodationModel.slx), incorporating all the features discussed in the paper. The Simulink model has two inputs: (1) demand, and (2) whether or not the eye is viewing through a pinhole. It has one output: accommodation.

To prevent the closed-loop instabilities from catastrophically destabilizing the response, we clip this non-predictive signal at a low value, set to 0.15D in our model (i.e. signals larger than 0.15D in magnitude are set to ±0.15D depending on their sign). This saturating value is chosen simply because it gives a reasonable match to empirical results. It is low enough to ensure that the signal does not change the behavior of the model in response to large changes in defocus. However, it is large enough that the signal still produces a visible high-frequency peak in the power spectrum of closed-loop microfluctuations and a high-frequency resonance in the response to low-amplitude sinusoids (see Results).

This non-predictive saturating signal has other interesting effects on accommodation. Notably, it facilitates a rapid response to small step stimuli, because non-predictive proportional signals tend to react faster than predictive integral signals. For example, suppose demand suddenly increases by 0.1D, causing an 0.1D step-change in defocus. The non-predictive proportional control signal, with unit gain, requests the full 0.1D increase in accommodation. The fast integrator begins responding at the same time, but due to its integral nature, its response ramps up more gradually. Furthermore, because the non-predictive proportional signal uses the current sensed defocus, rather than the predicted future defocus, it stays requesting the full 0.1D for at least 0.3s, until the sensorimotor latency has elapsed and the ocular plant starts to respond and thus reduces the sensed defocus. In contrast, input to the fast integrator is estimated future defocus, which begins to fall immediately based on the requested change to accommodation (the predictive control system assumes that demand will stay at the new value, but it predicts that defocus will fall because of the predicted accommodative response). So, the input to the fast integrator begins to fall immediately from its initial peak of 0.1D, whereas the input to the proportional controller stays at 0.1D until the sensorimotor latency has elapsed. Thus for small step-changes in defocus, the non-predictive proportional signal enables a larger, faster response. However, the saturation means that its effect is limited to small changes, with the predictive-integral control dominating the response to large changes. Dynamics of larger step responses are controlled with a pulse signal (Bharadwaj & Schor, 2005, 2006; Schor & Bharadwaj, 2004, 2006) that will be added to this model in a subsequent paper.

#### Depth of focus

In principle, changes in defocus that are so small they produce no detectable change in the retinal image, given the eye’s optics, cannot drive accommodation. The smallest change in defocus which produces a detectable change in accommodation is referred to as objective depth of focus. This is typically much smaller than the subjective depth of focus, i.e. the smallest change in defocus which produces a perceptible change in image quality (Kotulak & Schor, 1986a; Udlam et al., 1968; Yao et al., 2010). Depth of focus is often modelled as a deadzone (e.g. (Khosroyani & Hung, 2002; Schor, 1979b)): the defocus signal is set to zero unless it exceeds some threshold value corresponding to the objective depth of focus, say 0.2D. In such models, the deadzone contributes to lags and leads of accommodation, since the error signal vanishes once accommodation comes within the deadzone. However, this approach has a number of drawbacks:

i. It can result in unrealistic jumps, where a small change in demand pushes the defocus above the threshold and thus elicits a disproportionately large response.
ii. It produces a hysteresis effect, whereby accommodative lead and lag can depend on how the demand is approached. For example, with a threshold of 0.2D, if the demand steps up from 1D to 2D, the effective defocus becomes zero once accommodation reaches 1.8D, so we get a lag. But if demand steps down from 3D to 2D, effective defocus becomes zero once accommodation reaches 2.2D, so we get a lead. This hysteresis is not typically observed, except with extremely blurred images (Heath, 1956a).
iii. It reduces the gain of the response to low-amplitude oscillations. For example, consider a slow oscillation ranging between 1D and 3D. Assume for simplicity that the closed-loop gain of the system is 1, so that in the absence of a deadzone, the response would track demand exactly. With a deadzone clipped at 0.2D, the response would range from 1.2D to 2.8D, reducing the gain to 0.8. With a lower-amplitude oscillation where demand ranged from 1.5D to 2.5D, the response would range from 1.7D to 2.3D, making the gain 0.6. With a still lower-amplitude demand ranging from 1.7D to 2.3D, response would range from 1.9D to 2.1D, making the gain 0.3. Yet this decrease in gain with decreasing amplitude is not observed. In fact, accommodative gain tends to be smallest for high amplitudes, not for low amplitudes (Stark et al., 1965, p. 196).

Furthermore, recent evidence has undermined the experimental support for the notion of a deadzone. The accommodative system produces measurable responses to small amounts of defocus which do not introduce perceptible blur (Kotulak & Schor, 1986a; Yao et al., 2010), while the measured accommodative lags and leads may in fact maximize image quality rather than reflecting a deadzone (Labhishetty et al., 2021). For all these reasons, we have chosen not to include a defocus deadzone in our model. The objective depth of focus is adequately accounted for here by the white noise we have added to the defocus signal, which effectively swamps small changes. A more complete model would of course compute defocus from the retinal images, and thus take into account that small changes in defocus are hard to detect (Labhishetty et al., 2021).

### Simulink implementation and summary of the model

Figure 10 shows the complete model as it appears in our Matlab Simulink implementation, incorporating all the elements discussed above. The Simulink model has two inputs: (1) “demand”, accommodative demand in diopters, and (2) “pinhole”, a binary signal which conveys whether the eye is currently viewing through a pinhole or not. If pinholes are present, the defocus signal is set to zero at the block labelled “Apply Pinhole”; otherwise it is set to the optical defocus, i.e. demand minus accommodation. The defocus signal has white noise added to it and is delayed by the sensory latency before reaching the “brain” module.

Here, four signals are combined to produce a neural signal which is delayed by the motor latency before reaching the ocular plant. From top to bottom, these four signals are: (1) the constant bias signal, which sets the rest focus; (2) the proportional signal, which is simply the noisy defocus signal clipped at ±0.15D; (3) the signal from the fast integrator, which is driven by the estimated future defocus; (4) the signal from the slow integrator, which is driven by the fast integrator. One final detail not mentioned so far is that the neural signal is thresholded at zero to ensure it is positive. This is visible in the diagram as the “saturation” block on the far right, immediately after the four signals are combined. This accounts for the fact that the ciliary muscle can only be commanded to contract, making the lens more convergent, or allowed to relax. Negative values would effectively command the ocular lens to adopt a divergent form, which is physically impossible.

As well as being sent down cranial nerve III to the eye, an efference copy of the neural signal is directed to a virtual plant within the brain, which predicts the future accommodation. This in turn is used to estimate the future defocus which drives the fast integrator. For completeness, we have included a block labelled “Demand Predictor”, although in the current instantiation of the model, this simply passes its input through unchanged.

#### Simulation details

The next section shows simulation results for sine and step stimuli with this model. All simulations were run in Simulink, Matlab R2020b, with a variable-step solver, automatic solver selection and the default settings (relative tolerance 0.001 and max/min/initial step size and absolute tolerance all set to “auto”). For plotting, we interpolated the output to obtain results every millisecond. Note that this can give the impression of greater variability than in some empirical results where accommodation may be measured at a much lower rate, e.g. 50Hz. To obtain the velocity traces shown in Figure 16, we took the difference between successive accommodation values to obtain the change per millisecond, then smoothed this within a moving window of 10ms.

To obtain the model gain and phase in response to sinusoidal oscillations in demand, we ran the model for 25 cycles of the specified frequency, then fitted a sinewave to the results using Matlab’s Curve Fitting Toolbox. We fixed the frequency of the sinewave to the frequency of the stimulus and fitted the three free parameters baseline, amplitude and phase (see code in Run_Sine.m). The amplitude and phase of the response were taken to be those of the fitted sinewave.

The simulation shows onset transients at its start point, as the integrators settle. In all cases, we therefore discarded the first few seconds of simulation time in order to exclude these transients.

## Results

The different elements of this model were motivated by different observations – the gain and phase to sinusoids; adaptation; power spectra of microfluctuations so on. Components such as the fast and slow integrator and the virtual plant have been proposed before for the accommodation step response (Schor & Bharadwaj, 2005), but to our knowledge never tested in combination for pursuit sinusoidal tracking (Schor & Kotulak, 1986) or adaptation (Schor, 1979b), or with white noise and the feeding through of a clipped signal proportional to the current defocus. This combination is thus a novel contribution of this paper. We now demonstrate that this unified model can reproduce each of the observations that motivated its different components.

### Response to sinusoidal demand

Figure 11 shows the gain and phase of the model (heavy black line), compared with results from human subjects digitised from (Kruger & Pola, 1986; Ohtsuka & Sawa, 1997). This is of course similar to results already shown in Figure 7, but whereas those curves were obtained from mathematical formulae for a leaky integrator in a predictive control system, Figure 11 is obtained via Simulink simulation of the full four-signal model with noise. There is reasonable agreement in gain (Figure 11AB); both humans and model are low-pass. The main quantitative disagreement is that the “knee”, where the gain drops rapidly, typically occurs around 0.4Hz in humans and slightly later, around 0.6Hz, in the model. There is also good agreement in phase (Figure 11C). For comparison, the dashed black line shows the phase which would be obtained for a model with perfect demand prediction. This can be obtained from the phase of our model with “no change” demand prediction by subtracting the sensorimotor latency: ϕ_perfect_ = ϕ_nochange_ – 360*fT_sens_*. The phase function of most human subjects agrees better with that of the no-change model rather than the perfect model, suggesting that these subjects had little ability to predict the oscillatory demand.

**Figure 11.**
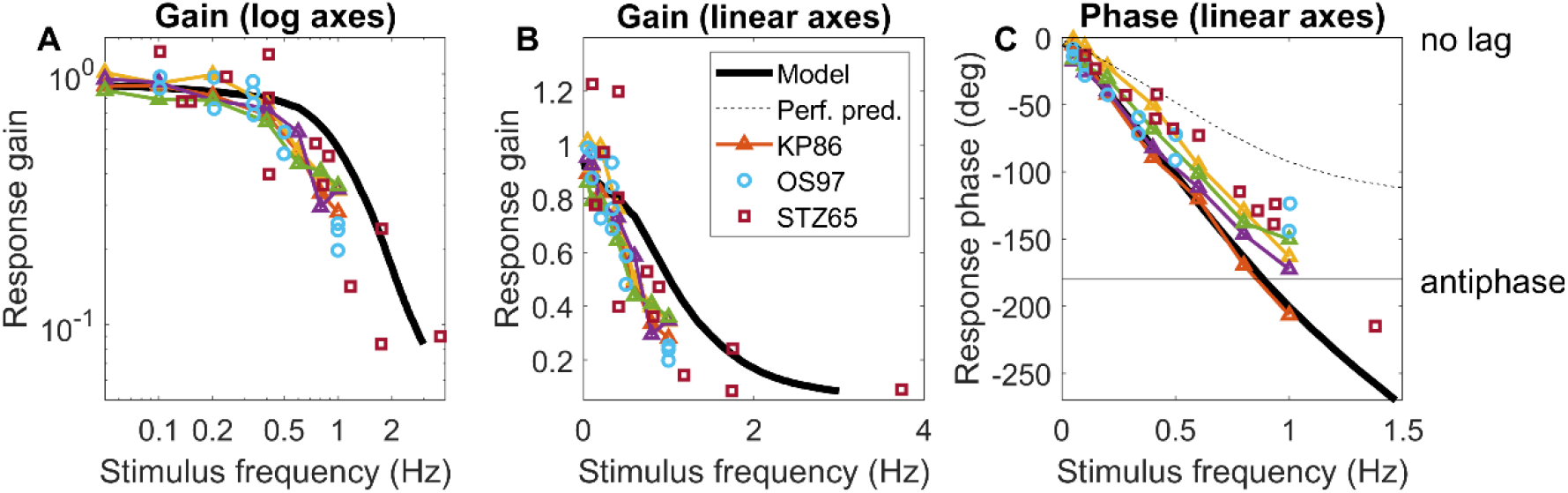
Gain and phase of the model response to sinusoidal demand, compared to empirical results. A,B: Gain plotted on linear and log axes. C: Phase plotted on linear axes. The heavy black line is the response of the model in Figure 10 with the parameters given in Table 2. The dashed black phase line shows the phase which would be obtained by a model capable of perfectly predicting the sinusoidal oscillation in demand. Triangles show empirical results for four human subjects, digitised from Kruger and Pola (Kruger & Pola, 1986), using the data with white light and defocus cue only. Circles are for a further four subjects, digitised from Ohtsuka and Sawa (Ohtsuka & Sawa, 1997), using only their control subjects. In (Kruger & Pola, 1986) and in the model, the demand oscillated between 1D and 3D, i.e. the amplitude of the sinusoid was 1D and its mean value was 2D. In (Ohtsuka & Sawa, 1997), the amplitude was 1.5D and its mean value is not stated. Code to generate this figure is in Fig_CompareGainPhase.m. Run_Sine.m must be run first to generate the model data.

Figure 11 was for sinusoidal demand oscillations with an amplitude of 1D. Of course, the gain and phase of a linear system are independent of amplitude. However, our model is nonlinear due to the saturation of the non-predictive proportional signal. Figure 12 shows the gain and phase in the same format as Figure 11, but for different amplitudes of oscillation around a 2D baseline. The green lines are for the 1D amplitude shown in Figure 11, but for lower amplitudes the gain and phase start to deviate significantly from these results. Most strikingly, there is a resonance at 1.2Hz where the gain actually goes above 1 for the smallest oscillations (±0.1D). This represents the instability caused by the non-predictive proportional signal. Since this signal is clipped at ±0.15D, it has a significant effect only for low-amplitude oscillations.

**Figure 12.**
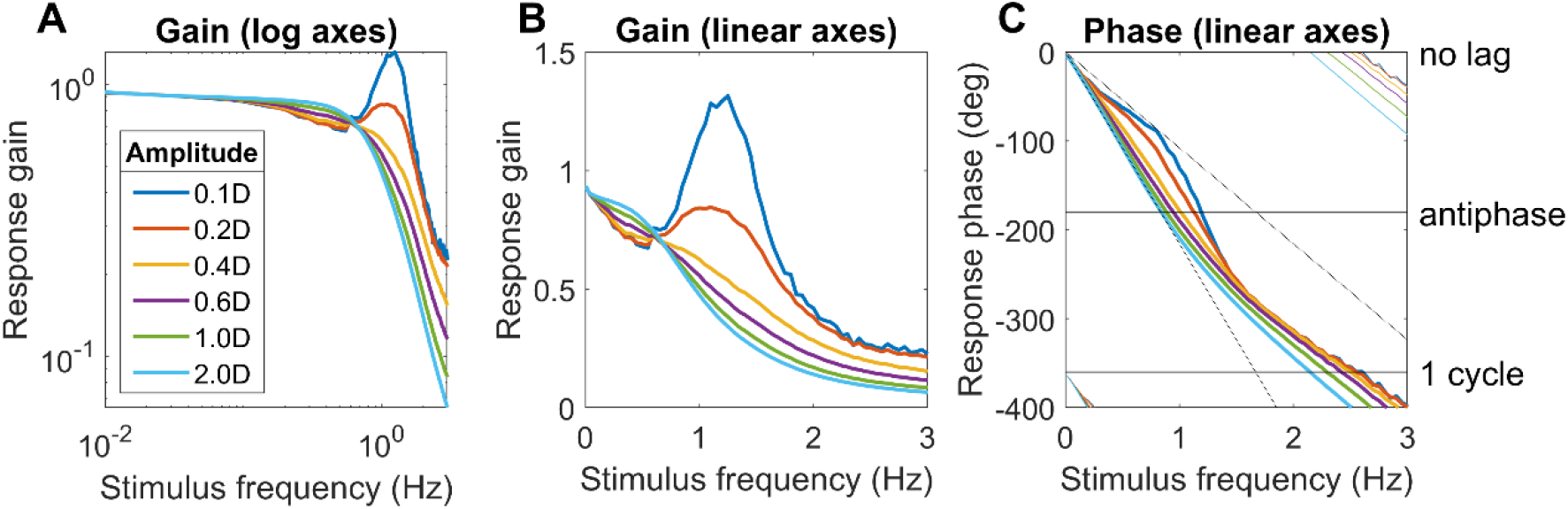
Model gain and phase as a function of amplitude. The green curves (1D) are what was shown in Figure 11, but we see that the behaviour at low amplitudes is quite different, with a resonance at 1.2Hz. Code to generate this figure is in Fig_Sine.m. Run_Sine.m must be run first to generate the model data.

This effect is qualitatively in agreement with the low-frequency resonance reported by Stark et al. (1965), which led them to conclude that human accommodative control must include a nonlinearity. Digitized data from Stark et al. (1965) is replotted in Figure 13, along with the response of the model. The model does not reproduce the strong dip in gain at 0.8Hz for an amplitude of 0.3D (blue point in Figure 13A), but apart from that, the agreement is quite good. In particular, it accounts for the key observation that gain is quite high, around 0.5, for 0.3D-amplitude oscillations at around 2Hz, whereas gain is much lower, around 0.1, for higher amplitude oscillations at this frequency.

**Figure 13.**
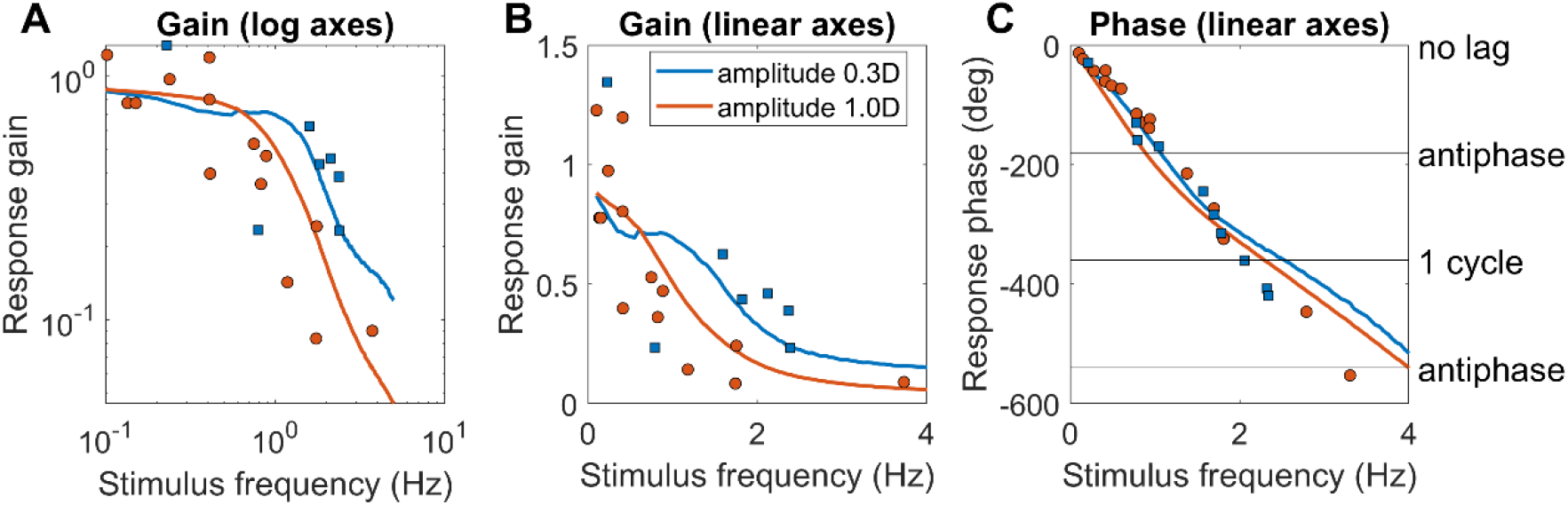
Symbols are digitised data from Figure 3 of Stark, Takahashi & Zames (Stark et al., 1965). These are measured gain and phase for one subject, for amplitudes of 0.3D (blue) and 1D (orange). The curves are model gains and phase for these amplitudes, about a baseline of 2D. Code to generate this figure is in Fig_StarkTakahashiZames.m. Run_StarkTakahashiZames.m must be run first to generate the model data.

### Limiting tracking frequency

When asking what are the fastest changes that can be tracked by accommodation, it is important to consider phase as well as gain. Figure 12C and Figure 13C showed that the phase of the response relative to demand increases with frequency, reaching 180 degrees at a frequency of around 1Hz. When this occurs, the demand and response are in antiphase, and the error is greater than the stimulus. Interestingly, if the response gain were zero, then the error for the 180 deg phase delay would be smaller than if the gain were 1.0. It is therefore of interest to ask how the gain and phase changes affect the model’s defocus error for demand oscillations of different amplitude and frequency. We quantify this using the mean absolute defocus error. The defocus error is the difference between demand and accommodation at any time; absolute defocus error is the rectified version of this waveform, and mean absolute defocus is the average value of this over time: 〈|*d*(*t*) – *a*(*t*)|〉, where *d*(*t*) = *D_mean_* + *D_amp_*(sin 2*πft*).

The heavy curves in Figure 14A show how mean absolute defocus error varies with amplitude and frequency of sinusoidal demand. In each case, the peak error is just below 1Hz, when the response is 180° out of phase with the demand (Figure 12C). The error increases with demand amplitude, even though for frequencies below the peak, the gain (i.e. the ratio of response to demand) is closer to 1 for larger amplitudes (Figure 12AB).

**Figure 14.**
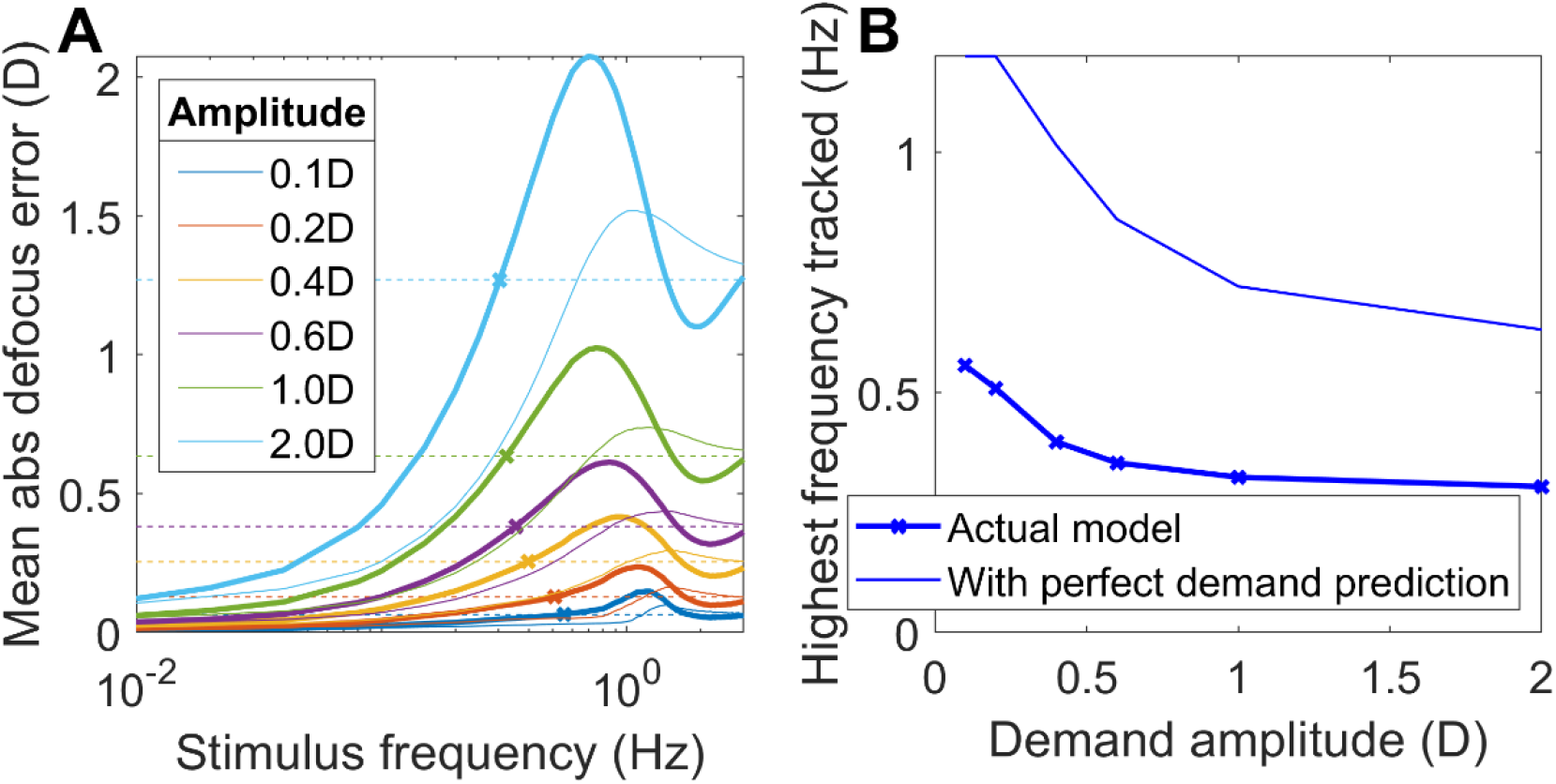
(A) Mean absolute defocus error for sinusoidal demand oscillations of different frequencies and amplitudes about a 2D baseline. The heavy curves show |d(t)-a(t)| for our model with its observed gain and phase; the light curves are those inferred for a model with perfect demand prediction. The dashed lines show the expected high-frequency limit, i.e. the mean defocus error if the demand oscillated but the response stayed at the steady-state value elicited by the mean demand, and the crosses indicate where this is first less than the error with tracking. The crosses mark where this crosses the mean defocus error. We take this as an indication of the highest frequency which can be successfully tracked at this amplitude. (B) Tracking frequency limit as a function of amplitude, for the actual model (heavy line, crosses) and for a model with perfect demand prediction (upper light line). Code to generate this figure is in Fig_Sine.m. Run_Sine.m must be run first to generate the model data.

The aim of accommodative control is to track demand so as to minimize defocus error, but the phase-delay means that for sufficiently high frequencies, this aim would be better achieved by simply keeping accommodation fixed at the mean demand, i.e. by having a response gain of 0, rather than attempting to track oscillations in demand about this baseline. The dashed lines in Figure 14A shows this *zero-gain tracking error*, i.e. the mean absolute defocus error which would be achieved if accommodation stayed at the steady-state value elicited by the mean demand (D_mean_=2D in this example). Because the amplitude of zero gain tracking error depends only on the input amplitude, the error is independent of temporal frequency of the sine input. Since the static accommodative lag is small, the zero-gain steady-state response is also close to 2D. So the mean zero-gain error is approximately the average value of |*D_amp_*(sin 2*πft*)|, or 2D_amp_/π, where D_amp_ is the amplitude of the demand oscillations about the 2D baseline.

We define the *limiting tracking frequency* to be the frequency at which the actual gain and phase-delay of the accommodative response produces the same error as would be achieved with zero gain. This is where the zero-gain tracking error is first equal to the actual error, marked with a cross x in Figure 15A. For frequencies lower than this limit, the oscillation in accommodative response is helpful, i.e. it tracks the oscillations in demand with a phase delay low enough to reduce the mean defocus error below the zero-gain tracking error. However for frequencies above the limit marked with a cross, the oscillatory response is out of phase and ends up making mean defocus error larger than if accommodation simply remained constant at the baseline value.

**Figure 15.**
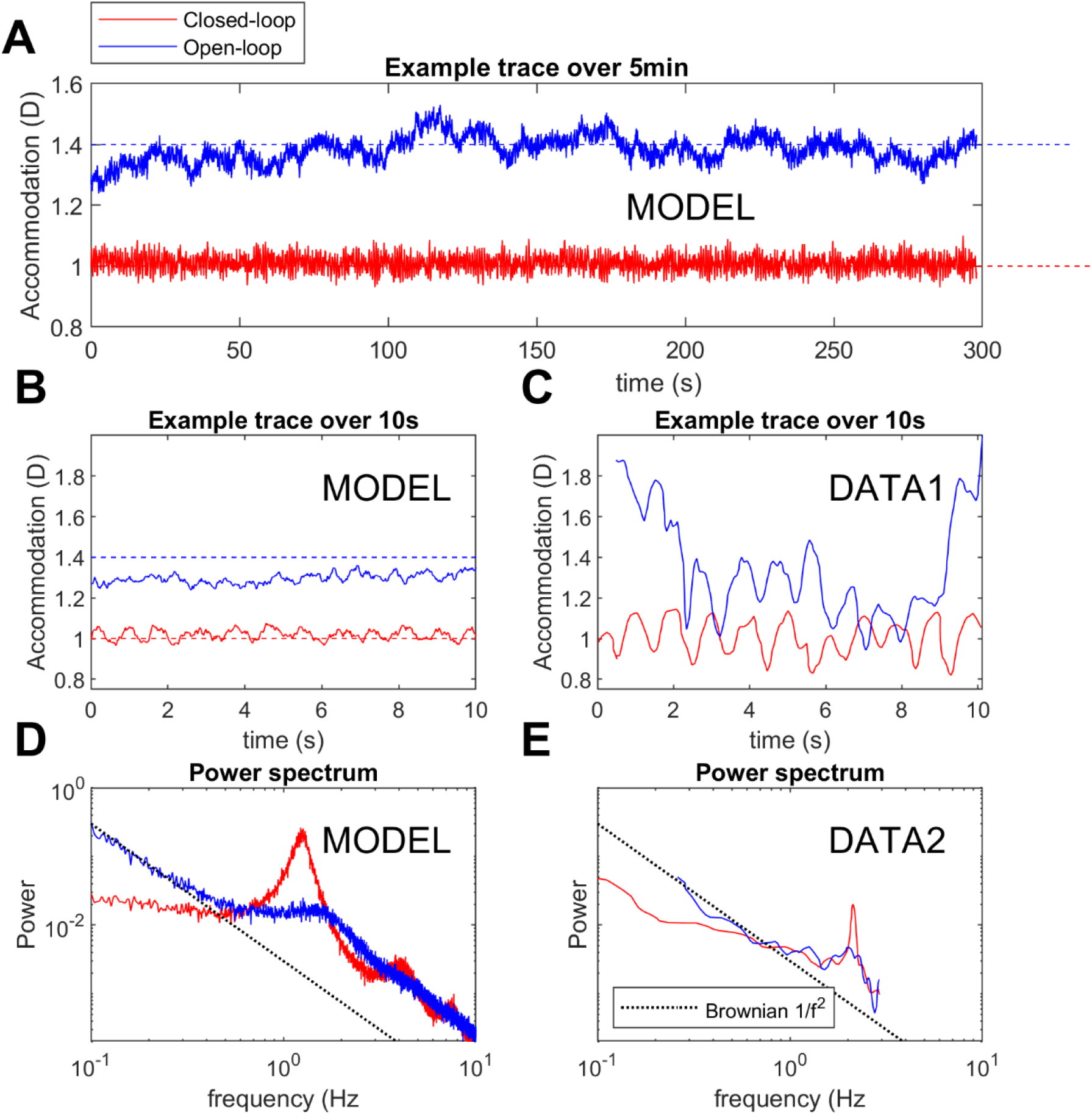
AB: Example accommodation traces in (red) closed-loop response to 1D and (blue) open-loop mode. Dashed horizontal lines show (red) the 1D demand and (blue) the 1.4D rest focus. A: trace over 5 minutes, to show slow fluctuations in open-loop response; B: 10s excerpt from A, to facilitate comparison with C: Example 10s trace recorded from a human observer, digitised from Fig 3 of Gray, Winn and Gilmartin (Gray et al., 1993a). The red trace is for a 5mm pupil; the blue trace is for viewing through pinholes of 0.5mm diameter. A scalebar but no accommodation values are provided in (Gray et al., 1993a), so the vertical position is arbitrary. To facilitate comparison with the model, we have set the mean value to 1D for the closed-loop and 1.4D for the open-loop trace. D: Power spectra of the closed- and open-loop response, obtained by averaging the Fourier power spectra of 50 traces like those in A, generated from simulations with different noise seeds. For comparison, a 1/f^2^ Brownian noise spectrum is drawn on with a black dashed line. E: Power spectra of closed- and open-loop responses for a human observer, digitised from Fig 5 of (Campbell et al., 1959b). This is labelled DATA2 to make clear that it is not the power spectrum of the trace shown in Figure 15C. No vertical axis scale was provided in (Campbell et al., 1959b), so we have scaled the spectrum so it best agrees with D. The red curve was recorded with a 7mm pupil and the blue curve with a 1mm effective entrance pupil. Code to generate this figure is in Fig_Noise.m; Run_Noise.m must be run first to generate the data.

Because of the nonlinearity represented by the saturating non-predictive proportional signal, this limiting-tracking frequency depends on amplitude, as shown in Figure 14B. For large-amplitude oscillations in demand, accommodation can track only up to around 0.4Hz. We saw above that the non-predictive proportional signal enables a more rapid response to small changes. This is shown in Figure 14B by the increase in limiting tracking frequency for low-amplitude oscillations.

Using the result that perfect demand prediction would reduce the phase by the sensorimotor latency, we can also infer what these curves would be for a model with perfect demand prediction but with the same plant and same leaky-integral controller. These are shown with the light curves in Figure 14AB. Perfect demand prediction does reduce the error and increase the limiting tracking frequency, but not dramatically, because of limits imposed by the time constants of the plant and of the fast integrator.

### Steady-state microfluctuations

We now turn to noise, and examine how well our model can account for accommodative microfluctuations. Figure 15A shows example closed- and open-loop accommodation traces recorded from the model over the course of 5 simulated minutes. The red trace is for closed-loop viewing of a stimulus at 1D (red dashed line). Accommodation thus fluctuates around a value a little over 1D, reflecting the accommodative lead for a stimulus nearer than the rest focus, here 1.4D. The fluctuations span a range of around 0.1D (±2SD). The SD is 0.03D, which is small compared to the SD of human microfluctuations (0.1-0.3D, (Charman & Heron, 1988, 2015; Gambra et al., 2009)). The power spectrum, Figure 15D, has a prominent peak at around 1.5Hz. This periodic structure is clearly visible in the 10s excerpt from the trace shown in Figure 15B.

The blue trace is for open-loop viewing, e.g. through pinholes. Now, the response wanders around the rest focus, 1.4D (dashed blue line). However, because the bias signal is constant rather than scaling with the difference between accommodation and rest focus, the excursions are much wider. This is visible in the power spectrum, Figure 15D, where the power continues to rise as frequency reduces.

Figure 15B shows a 10s excerpt from the trace in Figure 15A, for comparison with the example empirical data in Figure 15C, digitized from (Gray et al., 1993a). Although the amplitude of the microfluctuations is larger in the human observer, the same qualitative features are visible: closed-loop mode showing strong periodic structure at around 2Hz, open-loop mode showing much larger low-frequency fluctuations. Figure 15E shows the closed- and open-loop power spectra for a human observer, digitized from (Campbell et al., 1959b), for comparison with Figure 15D. Despite quantitative differences, they show the same qualitative features, notably a much larger peak for closed-loop.

The presence of this relatively large, 1-2Hz periodic component in the closed-loop microfluctuations may aid accommodative control, for example by “hunting” for the point of optimal focus (Kotulak & Schor, 1986c). Thus, this could be a reason why the postulated non-predictive proportional signal is beneficial for accommodative control.

### Response to step changes

When motivating the introduction of the saturating non-predictive proportional signal, i.e. a proportional controller responding to the current defocus signal (Figure 10), we discussed why it produces a larger, more rapid response to small changes in demand. We have already seen how this effect produces a higher gain for high-frequency low-amplitude oscillations (Figure 12) and thus the ability to track low-amplitude oscillations out to higher temporal frequencies than is possible for larger amplitudes (Figure 14). Similarly, the non-predictive proportional signal, clipped at ±0.15D, enables a faster response to small step changes in demand.

Figure 16 demonstrates this by comparing results from the full model (blue) with those from a model identical except that it lacks the non-predictive proportional signal (orange). To enable the effects to be seen clearly, noise is also turned off in this simulation. On the left, Figure 16AC, we plot the accommodation and velocity for a 0.5D increase in demand. The model with the non-predictive proportional signal responds more quickly. However, for the larger 2D step shown on the right (note different y-scales), the saturation of the non-predictive proportional signal at 0.15D limits its effect, and it makes barely any difference either to accommodation itself or to velocity. In fact, for large step changes like that shown in Figure 16BD, there appears to be a fifth signal, a nonlinear pulse triggered by sudden large changes in demand (Schor & Bharadwaj, 2004, 2006). The pulse accounts for the empirical observation that the peak acceleration of the response for step increases in demand is roughly independent of the step size, instead of scaling with step size as would occur for a linear system. While implementing the pulse is beyond the scope of this paper, we note that the non-predictive proportional signal already moves in the right direction by boosting the acceleration for small steps, and thus helping reduce the difference between acceleration for large and small steps. This could be another reason for the accommodative control system to include the postulated non-predictive proportional signal.

**Figure 16.**
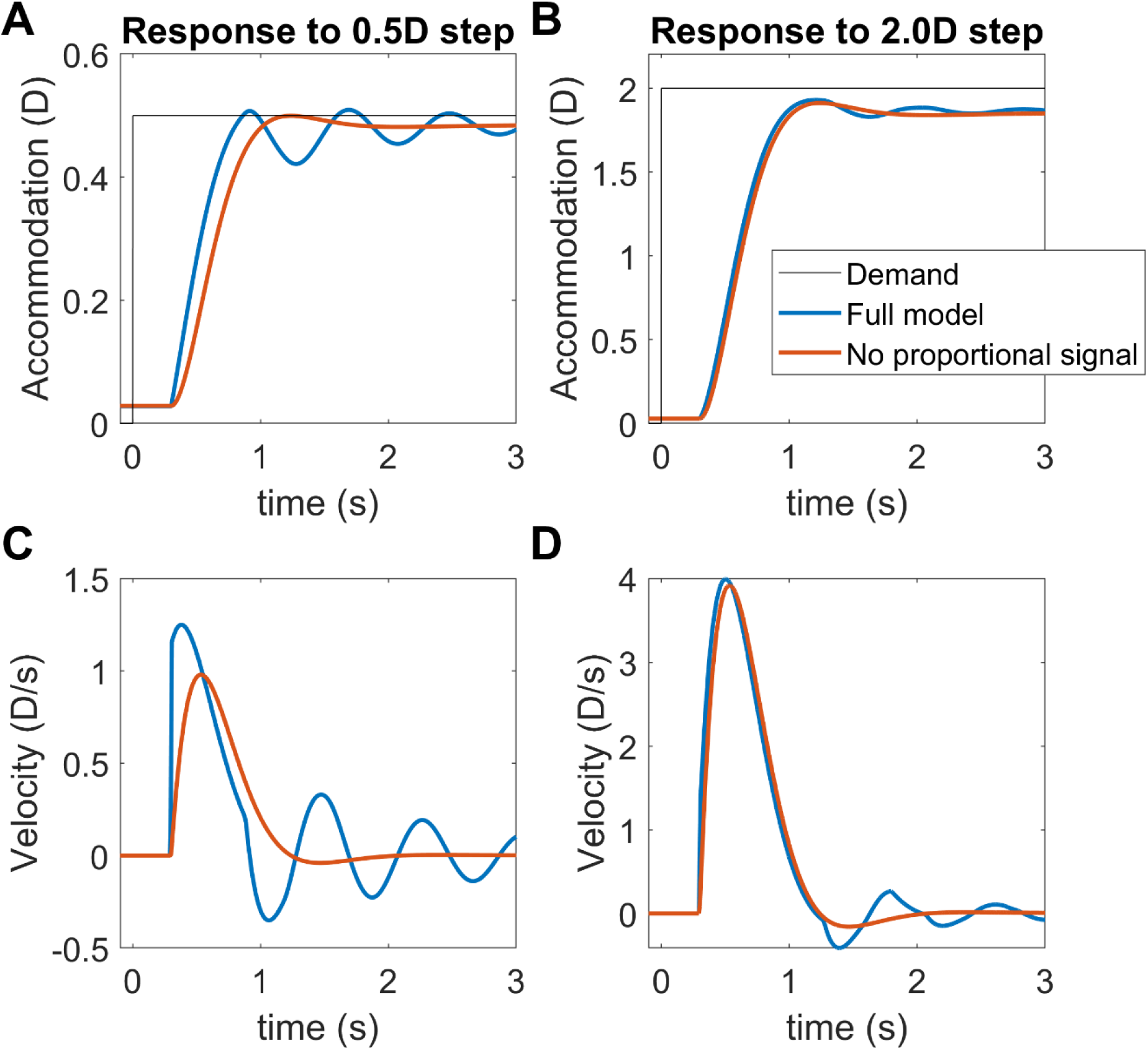
Noise-free accommodation (AB) and velocity (CD) for two different step increases in demand (AC: 0.5D, BD: 2D). The blue curve is for the usual model; the orange curve is for a similar model with no non-predictive proportional signal. To enable the effect to be seen clearly, noise has been turned off for this figure only. Also note that the response to the 2D step (BD) is included only to demonstrate the role of the non-predictive proportional signal. The model presented in this paper does not accurately capture the dynamics of the response to such large steps, since it does not include the pulse signal (see text). Code to generate this figure is in Fig_EffectOfPropSignal.m.

The blue curves in Figure 16 also show the ringing characteristic of non-predictive models, especially prominent relative to small step changes (Figure 16A). This instability is of course what we are exploiting to drive the high-frequency peak in the microfluctuations. Thus, our model predicts, probably wrongly, a transient increase in the amplitude of microfluctuations following small step changes in response.

### Adaptation

Next, we examine how the model adapts to accommodative demand to which it is exposed for more than a few tens of seconds. This was the motivation for postulating the slow integrator (Schor, 1979b; Schor et al., 1986). Its presence has not contributed to the results presented so far, other than to boost the gain for very slow oscillations. Now we see how it accounts for adaptation.

Figure 17 shows the time course of accommodation following the application of pinholes at t=0, shifting the system from closed-loop to open-loop demand. After the application of pinholes, accommodation eventually ends up at the rest focus, but how rapidly it does so depends on the demand before pinholes were applied. The model observer is initially adapted to 0D, then switches to viewing 3D for variable amounts of time as shown in the legend. The results show that after viewing one demand for at least two minutes, the observer adapts to it such that accommodation remains close to the adapted value for several minutes after pinholes have been applied (uppermost/red, lowermost/blue traces). Conversely, when the observer was exposed to different demands immediately before pinholes are applied (middle traces), they move much more rapidly to the rest focus.

**Figure 17.**
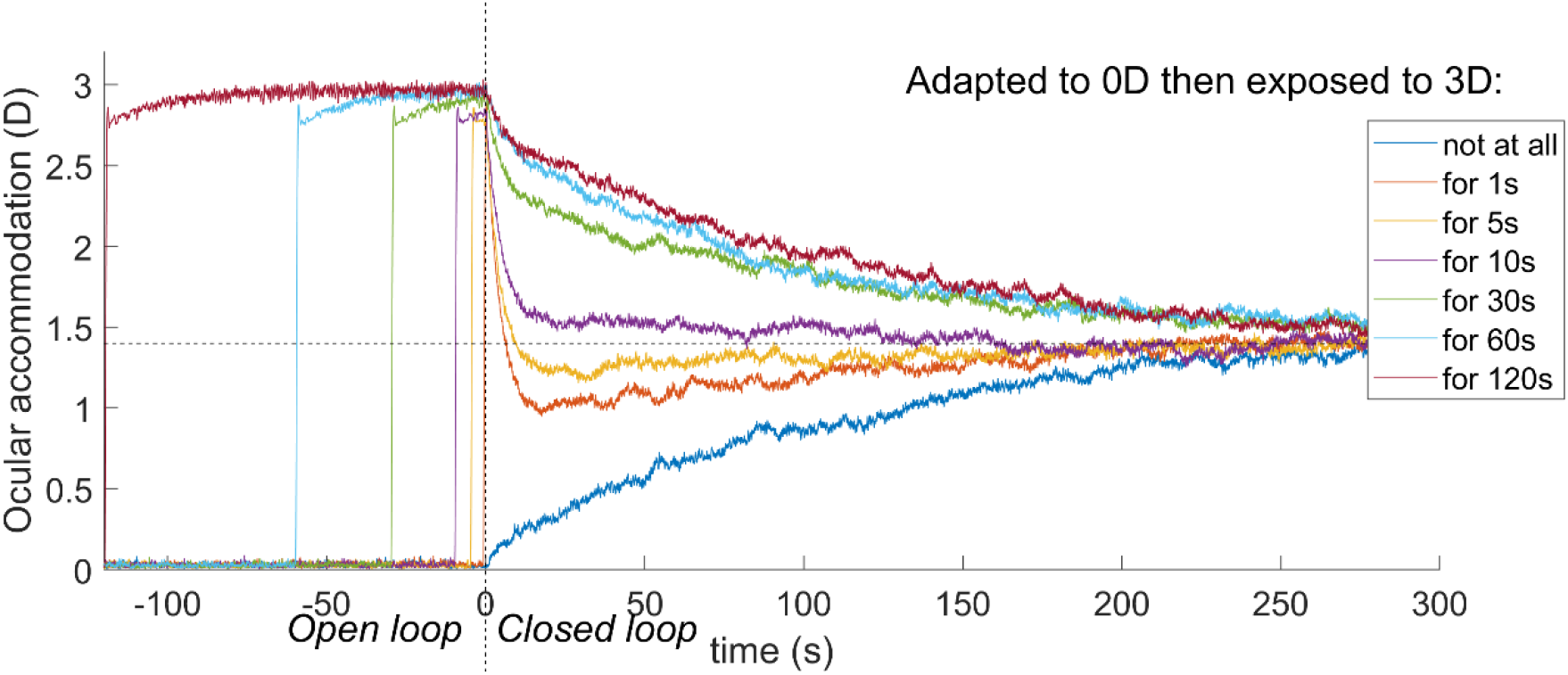
The model shows adaptation to demand, due to the slow integrator. The model observer is initially viewing an object at 0D, before then viewing an object at 3D for varying durations as shown in the legend. Pinholes are applied at t=0s, putting the system in open-loop mode. After long exposures, accommodation adapts to the demand, and moves only slowly to the rest focus; the adaptation affects the accommodation for many minutes after pinholes have been applied (e.g. dark blue curve: adapted to 0D, further than rest focus; red curve: adapted to 3D, closer than rest). Code to generate this figure is in Fig_Adaptation.m; Run_Adaptation.m must be run first to save the data in Results_Adaptation.mat.

Figure 18 shows a comparison with empirical data. Here, the observer was exposed to a demand of 2D for either 5s (blue) or 60s (orange) before moving to open-loop mode at t=0. The traces in Figure 18A are for a human observer (Schor et al., 1986); those in Figure 18B are from the model, with rest focus set to 0.4D (dashed line) in order to better match this observer. In both cases, following the 5s exposure to 2D, accommodation falls rapidly once the system enters open-loop mode, but following the 60s exposure, the decay is much slower.

**Figure 18.**
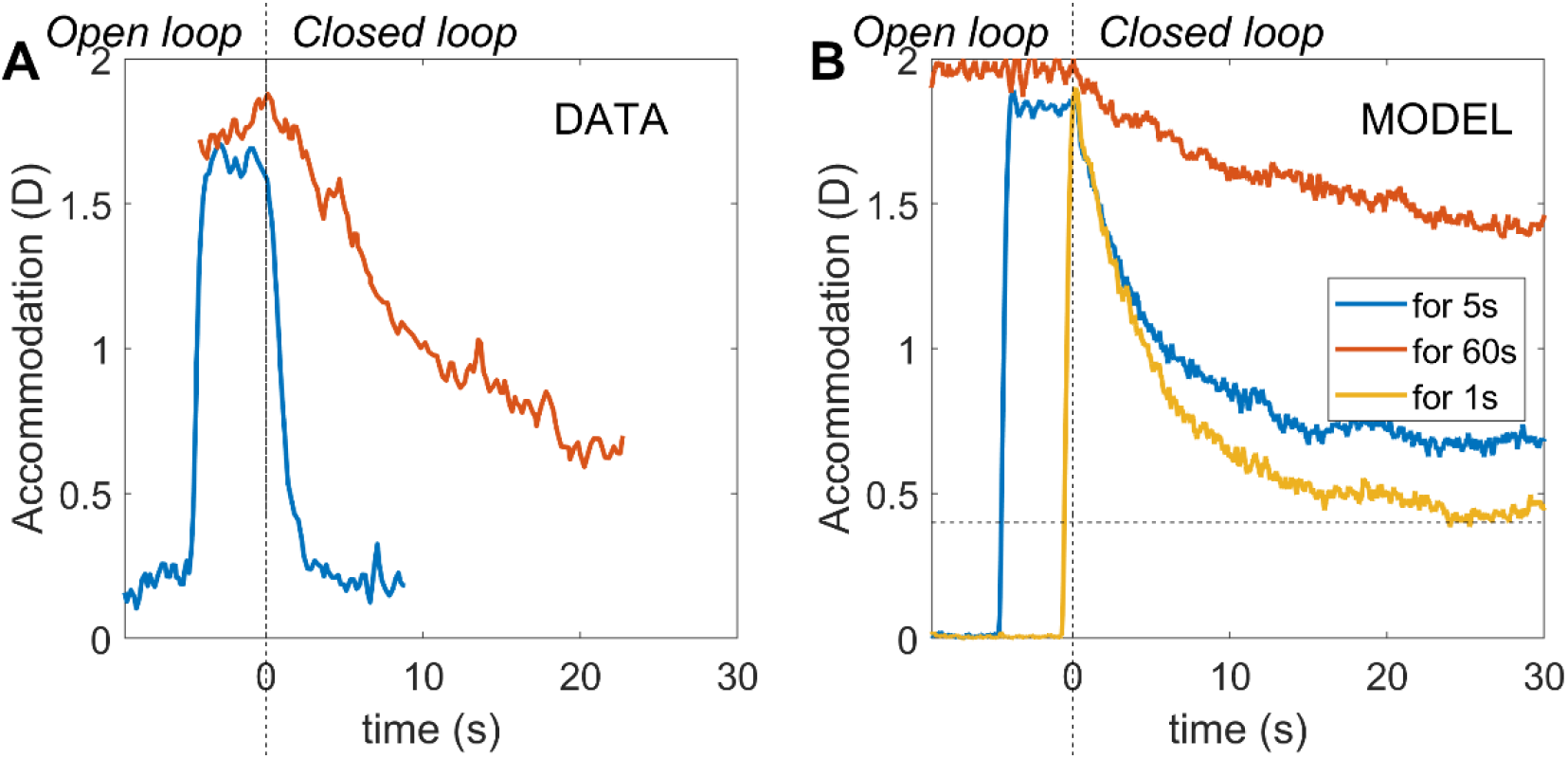
Comparison of the model (B) with data digitised from Schor, Kotulak & Tsuetaki (Schor et al., 1986) (Fig 2, empty field condition). As in Figure 17, pinholes are applied at t=0. Before then, the demand is at 0D for a long period, before moving to 2D for either 5s (blue) or 60s (orange); for the model, we also include 1s (yellow). In (Schor et al., 1986), a scalebar is provided, but absolute dioptre values are not available. The vertical position in the DATA panel is therefore arbitrary. However, since the open-loop condition decays by well over 1D from the closed-loop position adopted in response to a 2D demand, it seems clear that the rest focus for this observer was well below 1.4D. For this comparison, therefore, the rest focus of the model has been set to 0.4D (dashed line) in this figure only. Code to generate this figure is in Fig_SchorKotulakTsuetaki.m.

### Steady state error

Finally, Figure 19 shows the model’s steady-state error. As discussed (Equation 14), this reflects both the fast and slow integrator. In the model, the additional gain provided by the slow integrator means that steady-state error eventually becomes extremely small. Figure 20 shows this process for an example step up to 2D. The error is zero at the resting focus but shows lag/lead on either side of this. The gain (response/demand) therefore becomes high as demand tends to zero.

**Figure 19.**
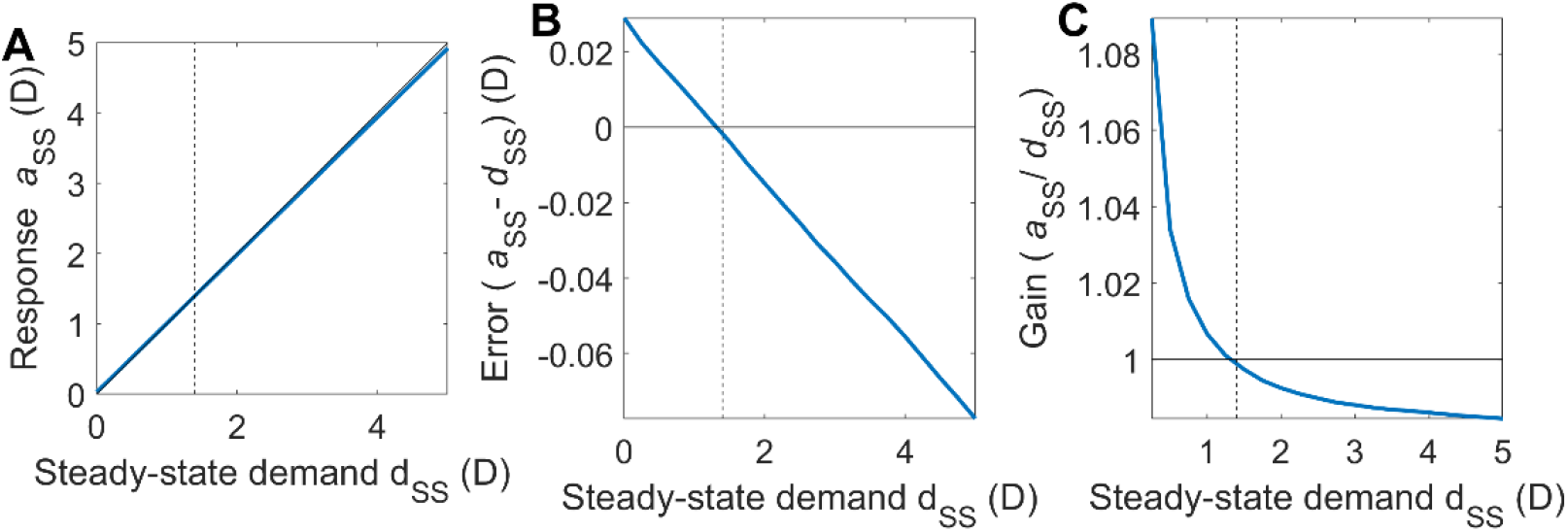
Steady-state response of the model. The model was run for 320s with a constant demand d_SS_ indicated by the value on the x-axis, and accommodation was averaged over the final 60s to obtain the steady-state response, a_SS_. (A) Input/output function, i.e. steady-state accommodation as a function of demand. (B) Steady-state error, i.e. difference between response and demand. For distant stimuli, this is positive (lead); for near, it is negative (lag). (C): Gain, i.e. ratio of response to demand. In each case, blue curves show the response of the model; solid black line indicates response equal to demand, and the dashed vertical line marks the rest focus, where this occurs. Code to generate this figure is in Fig_SteadyState.m; Run_SteadyState.m must be run first to save the data in Results_SteadyState.mat.

**Figure 20.**
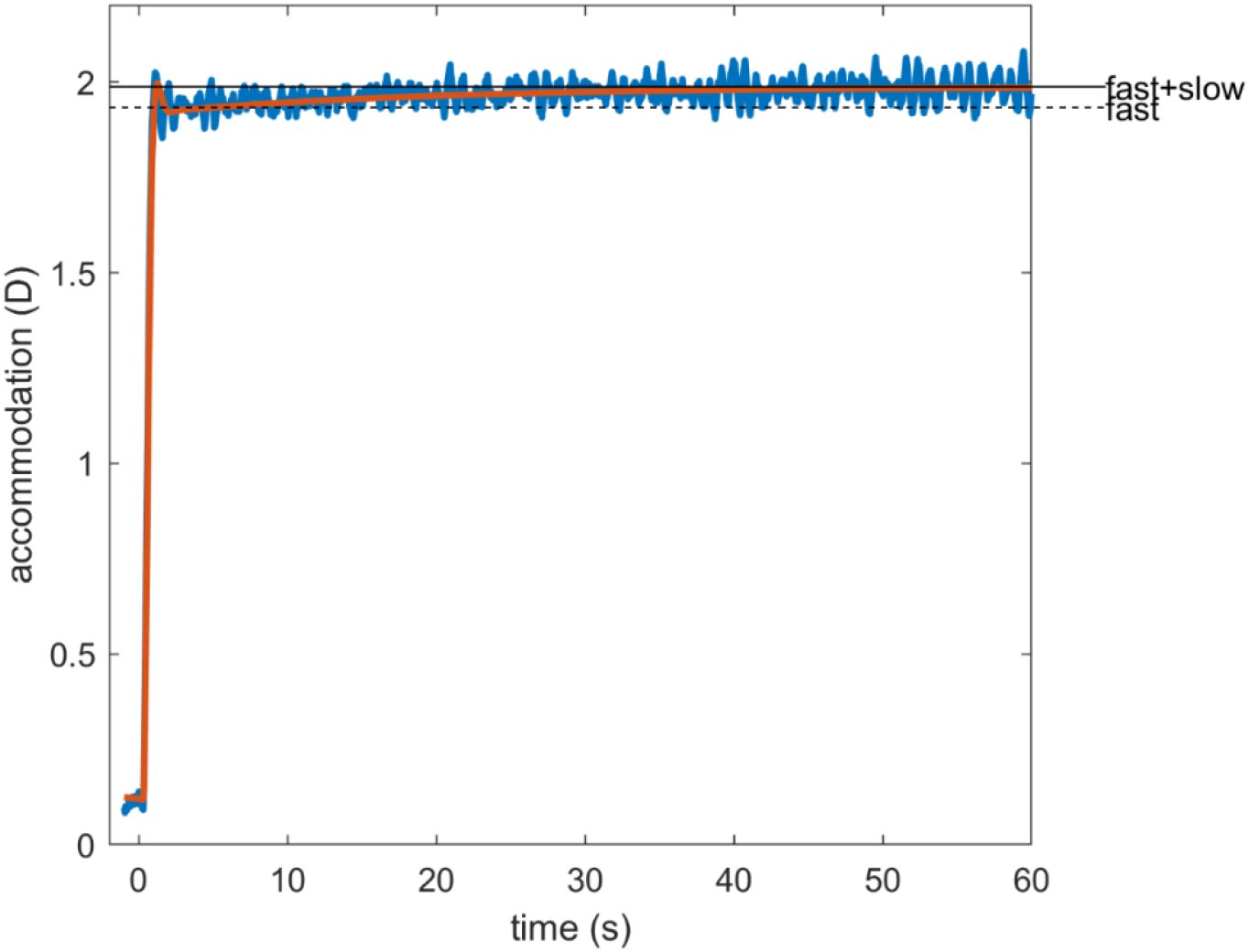
Model response to a step change in demand from 0D to 2D, showing the immediate rise to 89% of demand due to the fast integrator, and the subsequent slow rise to 98% demand due to the slow integrator. The blue trace is one example run from the full model; the superimposed orange line shows the response with no noise and no non-predictive proportional signal, in order to isolate the response due to the fast and slow integrators. Note that the dynamics of the immediate response to the step are not correct because they do not incorporate the pulse signal, but the point of this figure is to demonstrate the time-course following this immediate response. Code to generate this figure is in Fig_ExampleStep.m

## Discussion

In this paper, we have discussed the neural control of accommodation. We have provided a tutorial overview of the relevant control theory and key empirical observations. We have discussed the evidence for a predictive control system, i.e. one incorporating a forward model to predict the accommodative response in advance of the motor latency (Hung et al., 2002; Khosroyani & Hung, 2002; Schor & Bharadwaj, 2004). Similar models have also been proposed for vergence control (Erkelens, 2011; Hung et al., 1986; Zee & Levi, 1989) and saccades (Chen-Harris et al., 2008). Our analysis has led us to make the novel proposal that a saturating non-predictive proportional-control component may operate in parallel to the main predictive integrative-control feedback loop. This non-predictive proportional signal causes a high-frequency resonance in the closed-loop response, observed in the response to low-amplitude sinusoidal oscillations in demand. It amplifies noise within the system, explaining the high-frequency peak observed in closed-loop but not open-loop accommodation microfluctuations. It also speeds up the response to small, sudden changes in demand. Yet its saturation means that it does not destabilize the system as a whole, and that it becomes insignificant for large changes in demand.

We have implemented these ideas in a Simulink model, and are publishing this and all code along with the paper. Although most of the components of the model have been published before, we believe that this model is the first to incorporate realistic sensorimotor latencies, non-zero rest focus, noise *and* dual control by fast and slow integrators, as well as our novel use of a non-predictive proportional-control signal. Accordingly, it is able to account well for a wide range of empirical observations: the gain and phase of the response to sinusoidal oscillations in demand, including the puzzling high-frequency low-frequency resonance; the power spectrum of microfluctuations in closed-loop and open-loop modes, and the adaptation of accommodation to a steady stimulus.

### The four control signals: bias, fast, slow, non-predictive

In our model, accommodation is controlled by four separate signals (Figure 10), which offer different benefits. The constant bias signal sets the rest focus, to which the system returns in the absence of other stimulation (Figure 17). This may represent a typical demand, making it easier for the system to respond when stimulation restarts. The slow integrator means that the system tends to adapt to steady demand, perhaps reducing disruption if vision is briefly interrupted during sustained attention to one distance.

The fast integrator is the main workhorse of the feedback loop, enabling accommodation to respond rapidly yet smoothly to changes in demand (Figure 11, Figure 20). It is embedded within a predictive control system, incorporating a forward model to predict the effect of signals previously sent to the plant. This predictive control enables a smooth response and avoids ringing and instability. In principle, it can entirely remove delay due to the sensorimotor latency in a situation where demand can be predicted perfectly, as in a regular oscillation. However, it can slow the response to sudden and unpredictable changes in demand.

The fourth control signal can facilitate rapid responses in such situations (Figure 16). This signal is non-predictive: it is proportional to the currently sensed defocus, not the predicted future defocus. We originally rejected non-predictive control because it is prone to closed-loop resonances at particular frequencies. This is because the phase of the cycle where demand is high causes an increase in accommodation designed to null the defocus error, but – due to the latency – by the time the increase in accommodation has taken effect, the demand cycle has moved on to a phase where demand is low, and so the increase in accommodation in fact enhances the defocus error, causing a larger change in accommodation in the next cycle, and so on. In our model, we limit the destabilizing effect of this signal by making it saturate at low values. This ensures that it has little influence on accommodation in general, which remains dominated by the predictive integral control discussed above. However, the closed-loop resonance associated with non-predictive control remains detectable for small changes in demand. This amplifies noise within particular bandwidths, and means that the microfluctuations in the steady-state response show a peak at frequencies just over 1Hz, as observed. Opening the loop cuts the feedback pathway generating the resonance, explaining why this peak in the microfluctuation power spectrum is much less prominent in open-loop mode. The saturating proportional signal also accounts for the non-linear resonance observed when accommodation tracks low-amplitude – but not high-amplitude – sinusoidal oscillations in demand. However, an unrealistic feature of our model’s way of generating microfluctuations is that it predicts a transient increase in the amplitude of microfluctuations following small step-changes in demand – the ringing visible in Figure 16 – which has not been reported. The amplitude of microfluctuations in the model is also smaller than observed (Figure 15BC); this cannot be fixed simply by increasing the amplitude of the noise since that also changes the open- and closed-loop power spectra.

“Prediction” in the accommodation literature has often concentrated on predicting changes in demand (Krishnan et al., 1973; Stark, 1968). We believe it is helpful to draw a clear distinction between predicting one’s own accommodation, which is in principle possible perfectly with an efference copy and a forward model, and predicting demand, which is external and thus not always possible, for example when a fixated object moves suddenly. Predicting accommodation but simply using the current demand suffices to achieve closed-loop stability. The additional benefit of predicting future demand accurately is to avoid delay and thus avoid errors for rapidly changing stimuli. However, the low-pass characteristics of the plant and leaky-integral controller mean that the benefits of demand prediction are limited unless one also posits a different form of control.

### Deficits of the model

The model as currently implemented has many omissions and inadequacies, which must contribute to its imperfect ability to match the empirical data discussed in this paper. First, we do not consider control signals driven by inputs other than retinal defocus and bias (Heath, 1956b; Maddox, 1893). Notably, we do not include the crosslinks from and to the vergence system (Bharadwaj, 2005; Schor & Kotulak, 1986). We also do not consider other noise sources, such as heartbeat.

Second, this paper has nothing to say about how a signed estimate of defocus is obtained from the retinal image. This deficiency is perhaps especially important since our model assumes that visual feedback from the retinal images is the only feedback available to the accommodative control system. (Stretch receptors in the scleral spur base of the ciliary body could potentially also provide sensory feedback used in accommodative control (Tamm et al., 1994; Tamm & Lütjen-Drecoll, 1996), but at present nothing is known about whether or how this occurs, and it has not been included in any model of accommodative control.)

Perceptually, the threshold for detecting a change in focus which produces a detectable change in the image is higher for sharp than for blurring images: a focus change of 0.2D may be visible when the baseline defocus is 1.5D but not when the baseline is 0D (Campbell & Westheimer, 1958). It seems likely that such differences also affect the stimulus to accommodative control, but this is not taken into account in our model.

Related to this, we assume that the control system is attempting to minimize defocus, whereas in fact it is presumably attempting to maximize image quality. The accommodative lag and lead, which in our model is accounted for by the finite gain of the fast integrator, may effectively be an artefact of objective measurements of accommodation (Labhishetty et al., 2021). Recasting the control system so as to maximize a realistic measure of image quality rather than to minimize defocus could therefore profoundly alter the behavior of the model and lead to different conclusions about the nature of neural control. This would need to consider not only defocus but also higher-order aberrations such as spherical aberration, and should take into account pupil size. This would be computationally demanding to implement, and no published model of accommodative control has yet attempted it, but it must certainly be done in order to understand accommodative control in full.

The current model does not incorporate physical limits on accommodation, a non-zero far point or refractive error, nor do we consider how the system parameters may change with age (Bharadwaj & Schor, 2005; Schor & Bharadwaj, 2005), though these would be simple to add if required.

The model components are highly simplified. For example, the ocular plant is modelled as a linear-time-invariant leaky integrator with a fixed gain and time-constant, and the optical power is assumed to be proportional to the output of this integrator. A more accurate, yet usably simple, optical/biomechanical model of the relationship between ciliary muscle signal and optical power would be welcome (Wang et al., 2017).

The model developed here cannot account for the non-linear dynamics observed in response to large step changes. These have been accounted for previously with an additional “pulse” signal triggered by large step changes in accommodation (Schor & Bharadwaj, 2004, 2006), which temporarily overrides the error-driven signal, although non-linearities in the plant could also contribute. This of course means that the model presented here cannot accurately model the dynamics of the accommodative response to large step-changes, though it should remain valid for all the situations modelled in the Results (except Figure 16BD, included for illustrative purposes).

Finally, we have not attempted a realistic implementation of the demand-prediction model. There is some evidence that the brain can predict changing accommodative demand some time into the future, but we have here assumed it simply assumes demand will stay constant (Khosroyani & Hung, 2002). We hope to address some of these deficiencies in future work.

## Appendix

Here, we derive the total transfer function corresponding to the three types of linear models discussed in the text: (i) the non-predictive model and the predictive models with (ii) perfect and (iii) no-change prediction of demand. The bias signal due to the rest focus *a_RF_* is included as an inhomogeneous “forcing” term. We handle this by defining *A*(*s*) and *D*(*s*) to be the Laplace transforms of *a*(*t*)–*a*_RF_ and *d*(*t*)–*a*_RF_, respectively, where *a*(*t*) and *d*(*t*) are accommodation and demand as functions of time. In this way, we can effectively ignore *a_RF_* when obtaining the transfer functions.

### (i) Non-predictive model

The system diagram for this model is given in Figure 3. Reading around this circuit diagram, we see immediately that

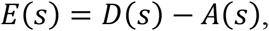

where E(s) is the Laplace transform of the defocus error signal, *d(t)-a(t)*. The input to the Controller block is *E*(*s*) exp(–*sT_sens_*), i.e. the defocus error signal after the sensory latency. The output from the Controller block is *C*(*s*)*E*(*s*) exp(–*sT_sens_*), where C(s) is the transfer function of the Controller. After accounting for the motor latency, the input to the ocular plant is *C*(*s*)*E*(*s*) exp(–*sT_lat_*). So, the output of the ocular plant, i.e. accommodation, is

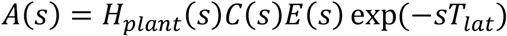

Substituting in for E(s), we obtain the closed-loop transfer function

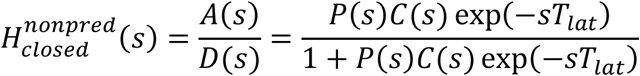

The gain and phase of the accommodative response to sinusoidal stimuli are the amplitude and phase of the complex number given by this closed-loop transfer function evaluated at s=jω=2πj*f*, *H_closed_*(2*πjf*). The closed-loop gain as a function of demand frequency is therefore

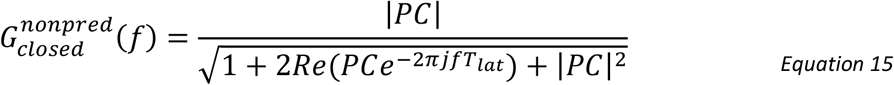

where the plant and controller transfer functions are similarly complex functions of frequency: *P=P(2πjf)*, C=*C(2πjf)*. The denominator contains oscillatory terms which mean that, even if *PC* is lowpass (i.e. a monotonically decreasing function of frequency), the denominator can be close to zero at particular frequencies and thus produce large resonances, for which the closed-loop gain exceeds 1. These manifest themselves as ringing or instability in the response to step changes in demand, and as gains>1 for sinusoidal oscillations in demand, which are not observed for large amplitudes.

With proportional control with unit gain (C=1), a sensorimotor latency of T_lat_=0.3s and the plant being a leaky integrator with τ_plant_=0.156s, Equation 15 has its first resonance at 1.2Hz where the closed-loop gain goes well above 1. This is ultimately responsible for the model’s high-frequency peak in microfluctuations (Figure 15) and the low-amplitude resonance in the response to sine-waves (Figure 12), although the precise behaviour also depends on the nonlinear clipping. The precise position of the first resonance depends on the gain of the proportional control, but only rather subtly. We therefore kept unit gain for simplicity.

We obtain the open-loop transfer function in the same way, but with the input to the Controller being D(s) instead of D(s)-A(s). This yields

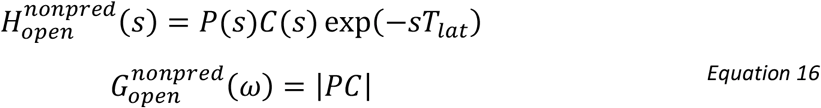

Thus, whether we use an integral or proportional controller in this non-predictive control system, the open-loop gain is purely low-pass, with no resonances. This means that adding our non-predictive proportional signal does not introduce any peaks to the power spectrum of open-loop microfluctuations.

#### Predictive models

The simplified system diagram for this model is given in Figure 5. As usual, we can ignore the bias signal if we express accommodation and demand relative to the rest focus. Reading around the circuit diagram, the demand signal is the input on the left; we represent this as usual in the Laplace domain by D(s). After passing through the sensory latency, it becomes *D*(*s*) exp(−*sT_sens_*), with the exponential being the Laplacian representation of a time delay (cf discussion of Equation 2). It then passes through the demand predictor, which attempts to predict the signal *T_lat_* = *T_sens_+T_mot_* into the future. If it did this perfectly, the output of the demand predictor would be *D*(*s*) exp(−*sT_sens_*) exp(−*sT_lat_*) = *D*(*s*) exp(+*sT_mot_*). To allow for the fact that demand is unlikely to be predicted perfectly, we will write the output as 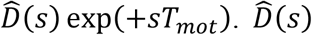 is the Laplace transform of the estimated future demand, again relative to the rest focus. That is, whereas *d*(*t*) is the actual demand at time *t*, 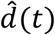 is the estimated demand at time *t*, as estimated at time (*t-T*_lat_).

Looking at the bottom of Figure 5, the output is accommodation, or A(s) in the Laplace domain. This is output after a motor latency T_mot_; thus the output of the “Plant” block in Figure 5 is *A*(*s*) exp(+*sT_mot_*).

Putting both these together, we see that the input to the Controller in Figure 5 is 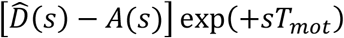. After multiplying this by the Controller and Plant transfer functions, we find that the output of the plant is 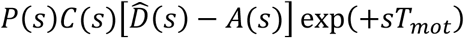. But we previously saw that the output of the plant is *A*(*s*) exp(+*sT_mot_*). Equating these, we see that

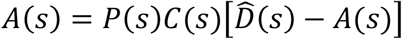

and thus that

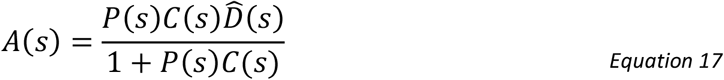

### (ii) Perfect demand predictor

In this idealized case, the demand predictor successfully outputs the future accommodative demand, so that 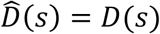 and the transfer function is

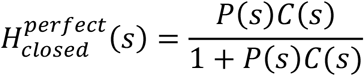

The closed-loop gain is therefore

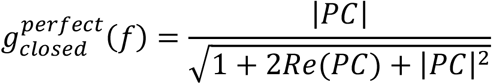

To obtain the open-loop transfer function, we replace D(s) with D(s)+A(s) in Equation 17, obtaining

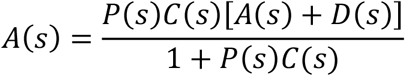

and thus

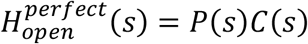

If demand prediction is perfect, the open-loop gain of the controller is independent of latency. For our situation where both the plant and controller are leaky integrators, the open-loop gain is lowpass, with no resonances.

### (iii) “No-change” demand predictor

In this opposite extreme, the demand predictor simply assumes that the future defocus after time T_lat_ will still be the same as the defocus it is receiving now:

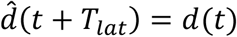

and thus

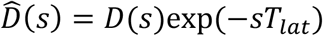

Hence

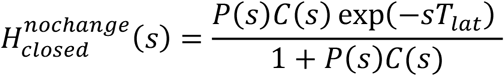

The closed-loop gain at any frequency *f* is therefore the same as for the perfect predictor, while the phase is reduced by 2π*f*T_lat_. In fact, the closed-loop gain would be the same for any demand predictor which accurately predicts demand any time at all into the future, even if, as here, that time is zero. Inaccurate predictions would of course change the closed-loop gain.

The open-loop gain does depend critically on demand prediction. With no-change prediction, replacing D(s) with D(s)+A(s) in Equation 17, yields

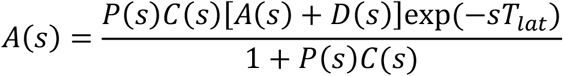

and thus

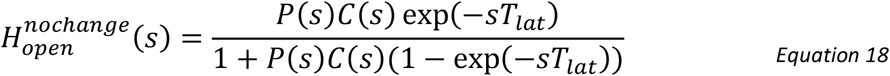

The presence of the oscillatory exp(–*sT_lat_*) term in the denominator can lead to local peaks in the gain at some frequencies. Thus with inaccurate no-change prediction, the system is prone to open-loop resonances due to the inner feedback loop via the efference copy. However, with our parameter values (Table 2), Equation 18 is a monotonically decreasing function of frequency. This ensures that we do not see local peaks in the power spectrum of open-loop microfluctuations (Figure 15).

#### The predictive model with leaky-integral control: a damped harmonic oscillator

For the case where the plant and the controller are both leaky integrators (Equation 9, Equation 10), and we neglect the other signals, the transfer function of the perfect-prediction model is

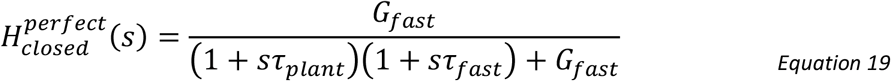

with *s* = 2π*jf*. This is the transfer function of a second-order damped oscillator. We can rewrite it in the standard form

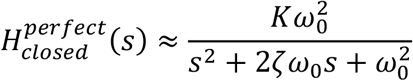

where K is the closed-loop gain:

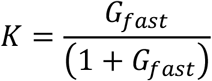

ω_0_ the natural angular frequency:

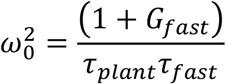

and ζ the damping coefficient:

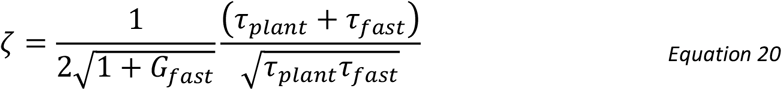

For perfect demand prediction, the phase at angular frequency ω is:

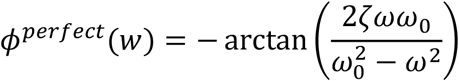

while for no-change prediction,

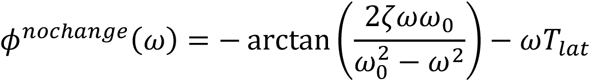

If ζ<1/√2, then the maximum gain occurs at the resonant angular frequency:

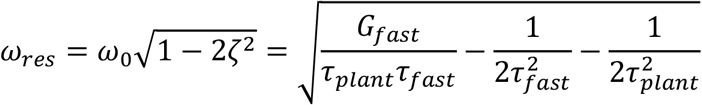

If ζ>1/√2, then the gain is maximum for f=0 and decreases monotonically with frequency. If ζ=1, the system is said to be critically damped.

As discussed in the text, to match the empirical gain of accommodation, ζ must exceed 1/√2, the minimum value for which gain decreases monotonically with frequency. Solving Equation 20, we find that

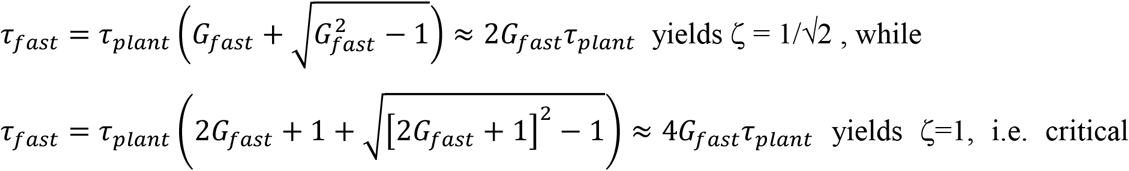

where the approximations hold since the gain G_fast_ has to be >>1, say at least 5, to avoid excessive lag. (Mathematically, there are two solutions, but the other one gives a very short time-constant for the controller, which in turn causes other problems such as open-loop resonances in the noise.)

##### The minimal-settling time solution

In the model presented here, we chose the “minimum settling time” solution which yields ζ = 1/√2:

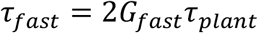

since this gave the best match to both gain and phase data. With this choice, since G_fast_>>1, the natural frequency is approximately

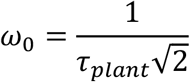

which with our value τ_plant_=0.156s corresponds to 0.72Hz.

For ζ=1/√2, the phase function is very close to linear out to ω=ω_0_. In this region, for perfect demand prediction

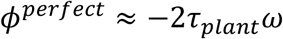

corresponding to an effective delay of *T*_delay_ = 2τ_plant_. Presumably coincidentally, this delay is very similar to the sensorimotor latency, although as we can see it arises from a completely different source. However, for frequencies beyond ~1Hz, the phase asymptotes to 180° (Figure 7).

For no-change prediction, the phase is approximately

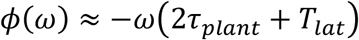

at low frequencies, corresponding to an effective delay of 2*τ_plant_* + *T_lat_*.

